# A Complex Containing HNRNPA2B1 and N^6^-methyladenosine Modified Transcripts Mediates Actions of Toxic Tau Oligomers

**DOI:** 10.1101/2020.12.03.409334

**Authors:** Lulu Jiang, Weiwei Lin, Cheng Zhang, Mamta Verma, Julian Kwan, Emily van Vliet, Peter E. A. Ash, Anna Lourdes Cruz, Samantha Boudeau, Brandon F. Maziuk, Shuwen Lei, Jaehyup Song, Victor E. Alvarez, Rakez Kayed, Nicholas Kanaan, Melissa E. Murray, Johnathan F. Crary, Leonard Petrucelli, Hu Li, Andrew Emili, Benjamin Wolozin

## Abstract

The microtubule associated protein tau oligomerizes in response to stress and disease, but the function of oligomeric tau (oTau) and the ultimate mechanisms of toxicity are unknown. To gain insights, we have now used Cry2-based optogenetics to induce tau oligomers (oTau-c) in neuronal cultures. oTau-c can seed tau aggregation and biochemical fractionates in a manner similar to oTau. Optical induction of oTau elicits a translational stress response that includes cytoplasmic translocation of the TIA1, abundant stress granules (SGs) and reduced protein synthesis. Proteomic analysis identifies HNRNPA2B1 as a principle target of oTau. Imaging and immunoprecipitation verify the HNRNPA2B1 association with endogenous oTau in neurons, animal models and human Alzheimer brain tissue. Mechanistic studies demonstrate that HNRNPA2B1 functions as a linker, connecting oTau with N^6^-methyladenosine modified transcripts (m^6^A). Knockdown of HNRNPA2B1 prevents oTau from associating with m^6^A, prevents oTau-induced reductions in protein synthesis and reduces oTau-induced toxicity. Finally, we show striking increases in m^6^A-oTau and -HNRNPA2B1 complexes in brains of Alzheimer subjects and P301S tau mice. These results reveal a novel complex containing oTau, HNRNPA2B1 and m^6^A that contributes to the integrated stress response of oTau.

**Highlights:**

1. Development of a powerful method combining optical induction of tau oligomerization with precision mass spectrometry to obtain time resolved evolution of protein interaction networks.
2. Demonstration of a tripartite complex that links tau oligomers with HNRNPA2B1 and N^6^-methyladenosine modified RNA in cytoplasmic stress granules.
3. Knockdown of HNRNPA2B 1 abrogates the interactions of oTau with N^6^-methyladenosine modified RNA, as well as inhibits oTau-mediated neurodegeneration.
4. Discovery that N^6^-methyladenosine modified RNA is significantly increased in the brains of P301S tau transgenic mice and in patients with Alzheimer’s disease.

## INTRODUCTION

The microtubule associated protein tau functions in health to stabilize microtubules. However, tau oligomerizes, becomes phosphorylated and accumulates in the somatodendritic arbor in response to stress and diseases such as Alzheimer’s disease (AD) and other tauopathies (Scheltens et al., 2016; Wang and Mandelkow, 2016). An immense body of literature describes the biochemical changes in tau occurring in response to stress and disease (Wang and Mandelkow, 2016), but left largely untouched are studies delineating the biological function of tau in stress and disease states. Increasing evidence suggests that these stress-induced changes in tau contribute to the stress response, but the molecular actors and mechanisms through which oligomeric tau (oTau) mediates its actions are unknown (Jiang et al., 2019; Lasagna-Reeves et al., 2011).

Previous studies of tau have relied on complex systems for inducing tau pathology. For instance, arsenite or other agents have been used to induce a stress response, which causes multiple forms of cellular injury and induces tau pathology. In vivo studies elicit tau pathology, but do so slowly and only in the context of aging, which obscures interpretation because of the large number of molecular events that occur in parallel. These complex conditions suggest that tau pathology is associated with strong changes in the biology of RNA binding proteins (RBPs) as well as the nuclear membrane (Apicco et al., 2018; Eftekharzadeh et al., 2018; Vanderweyde et al., 2016). Perhaps unsurprisingly, studies with induces stress responses show that pathological tau associates with RBPs, co-localizes with SGs and induces a translational stress response (Bishof et al., 2018; Chauderlier et al., 2018; Maziuk et al., 2018; Silva et al., 2019; Thompson et al., 2012; Vanderweyde et al., 2016). However, these studies all suffer because they are unable to distinguish between effects caused by tau itself from effects caused by the other pathways induced by concomitant stress. We now apply the light-reactive bacterial cytochrome 2 (Cry2) protein to generate oligomers of the tau-Cry2 chimeras oTau biology (Lamprecht, 2019; Shin et al., 2017), creating the ability to optically, selectively induce tau oligomerization in a temporally controlled manner.

We report that the optically induced tau-Cry2 oligomers (oTau-c) share many biochemical features in common with untagged oligomers, including an ability to seed tau aggregation in tau K18 sensor lines and a pattern of biochemical fractionation similar to that of oTau (Apicco et al., 2018; Sanders et al., 2014). Characterization of the tau protein:protein interaction (**PPI**) networks reveals a clear temporal evolution exhibiting greatly increased binding to RBPs. We identify HNRNPA2B1 as one of the strongest protein binding partners for oTau-c and oTau, as well as identify multiple other proteins genetically associated with AD that also bind to oTau. The role of HNRNPA2B1 as an indirect reader of N^6^-methyladenosine (m^6^A) tagged transcripts led us to investigate m^6^A in the biology of tau (Zaccara et al., 2019). We present the discovery that tau oligomerization induces striking cytoplasmic translocation of m^6^A, which co-localizes with HNRNPA2B1 and oTau. We further show that m^6^A levels are increased almost 4-fold in AD. Knockdown studies demonstrate that HNRNPA2B1 is required for localization of oTau with m^6^A as well as for the physiological actions of oTau, implicating HNRNPA2B1 as m^6^A reader (indirect) that links oTau to m^6^A labeled transcripts. These studies establish for the first time a complex containing oTau, HNRNPA2B1 and m^6^A that mediates actions oligomeric tau, and implicate m^6^A in the pathophysiology of AD.

## RESULTS

### Optogenetic Cry2 drives tau oligomerization in primary cortical neurons

To investigate the biology specifically related to tau oligomers, we created tau::mCherry::Cry2Olig (Tau::Cry2) and mCherry::Cry2Olig (control, mCherry::Cry2) chimeras which can inducibly trigger oligomerization through by fusing the Cry2Olig protein to the C-terminus of 4R1N WT tau::mCherry (or mCherry alone) in lentiviral expression vectors (**Figure 1A**). The constructs were transduced into primary cultures of hippocampal neurons with lentiviral vectors. Under basal conditions, Tau::Cry2 or mCherry::Cry2 constructs spread throughout the soma and dendrites. Tau::Cry2 associated with microtubules, consistent with the known homeostatic function of tau (Suppl. Fig. S1C). Exposure to low levels of 488λ blue light (200μW/cm^2^) induced rapid and robust oligomerization of both the tau::Cry2 or mCherry::Cry2 constructs (**Figure 1B, Suppl. Video 1, 2**). Tau::Cry2 was transduced in 60% of the neurons, but tau::Cry2 oligomerization occurred only in 30% of transduced neurons exhibiting strong tau::Cry2 expression. Neurons expressing low levels of Tau::Cry2 did not exhibit light induced oligomerization, while neurons exhibiting high expression exhibited robust oligomerization (**Suppl. Video 3**), which is consistent with prior observations of the Cry2 system (Shin et al., 2017). Transient exposure lasting <4 min induced granules for both types of constructs, with both sets of granules rapidly dispersing upon termination of the illumination (**Figure 1B**). However, with repeated or increasing duration of light exposure the oTau-c became more stable, showing no dispersion upon termination of the illumination (**Figure 1C, D**). In contrast, the mCherry::Cry2 oligomers remained rapidly reversible even after prolonged light exposures (**Figure 1B, D**).

**Figure 1.**
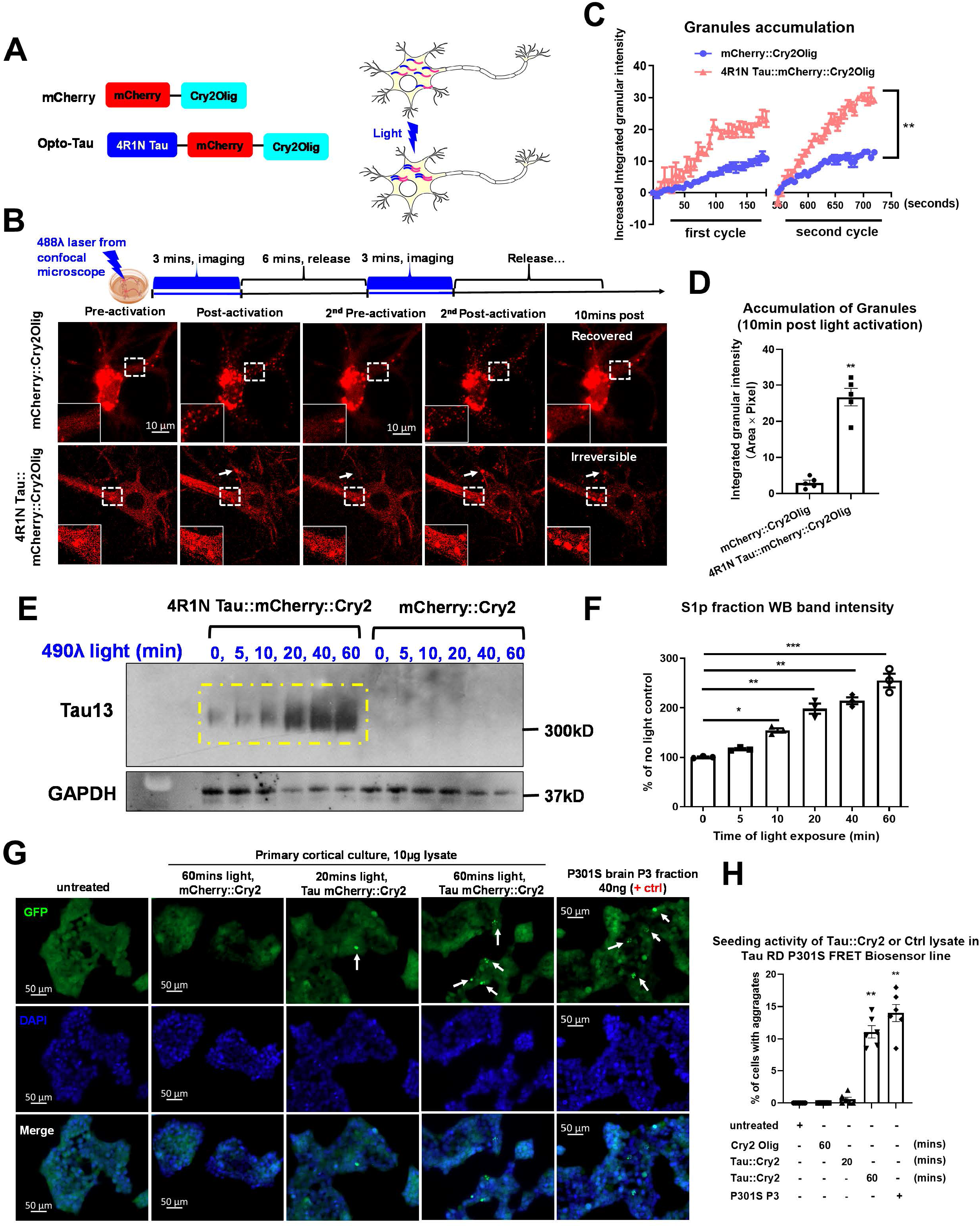
Optogenetic Cry2Olig drives tau oligomerization in primary cortical neurons. (A) Schematic diagram of the optogenetic platform. Full length 4R1N human tau fused to mCherry fluorescent protein and the Cry2Olig PHR domain. (B) Representative fluorescence images of blue light-activated assembly of mCherry::Cry2 or 4R1N Tau::Cry2 chimeras in living cells. The cells selected for live-imaging were at similar expression levels and activated under identical conditions. Scale bar, 20μm. (C) Quantification of rapid clustering of mCherry::Cry2 or 4R1N Tau::Cry2 chimeras in the first and second cycles of 488λ blue light activation in the experiments of (*B*), n=10. Error bars = SEM. Linear regression XY analyses \*\**p*<0.01. (D) Quantification of long-lasting oligomers at 10 mins after the termination of blue light illumination. n=10. Error bars = SEM. Unpaired T-test with Welch’s correction, two-tailed \*\**p*<0.01. (E) The Tau::Cry2 and mCherry::Cry2 transduced cultures were exposed to light for 0, 5, 10, 20, 40 and 60 min, respectively. The tau oligomer fraction was extracted from the total lysate by centrifugation. Tau was quantified in the pelleted fraction by immunoblot with the Tau13 antibody (recognizing all forms of tau). The resulting representative immunoblot demonstrates lightdependent increases of tau oligomers. (F) Quantification of the western blot in (E) with GAPDH as an internal control followed by being normalized to the “no light” negative control group. Data were collected from 3 independent experiments. Error bars = SEM. One-way ANOVA with Tukey’s multiple comparisons test was performed, \**p*<0.05, \*\**p*<0.01, \*\*\**p*<0.005. (G) Representative images showing the seeding activity of total lysate extracted from neurons expressing Tau::Cry2 exposed to 20 or 60 min light, respectively. The P3 fraction of fibrillar tau from PS19 mice (9 months old) was used as a positive control. Scale bar, 50μm. (H) Quantification of seeding induced by tau aggregates from the Tau::Cry2 or mCherry::Cry2 neurons. Data were collected from 6 independent experiments. Error bars = SEM. One-way ANOVA with Tukey’s multiple comparisons test was performed, \*\**p*<0.01.

The colocalization of Tau:Cry2 with microtubules was examined first to determine whether the Tau:Cry2 construct exhibited this association under basal conditions as is normally observed for tau. The distribution of Tau:Cry2 and microtubules were examined using immunofluorescence to characterize the fluorescent signal of anti-α-tubulin antibody labeling (green) and the Tau:Cry2:mCherry fluorescence (red) (**Suppl. Figure 1A**). Under basal conditions (no blue light stimulation) or conditions of short light exposure (5 min) Tau:Cry2 was observed to form filaments that colocalized with a-tubulin (**Suppl. Figure 1A**). With increasing duration of blue light exposure (>10 min), Tau:Cry2 formed round granules in neurons that did not colocalize with a-tubulin (**Suppl. Figure 1A**). Interestingly, this Tau:Cry2 granule formation was most abundant in the somatic region, while Tau:Cry2 in the dendritic fields largely retained its association with microtubules, as shown by the filamentous morphology and colocalization with a-tubulin (**Suppl. Figure 1A**). These results suggest that the Tau:Cry2 construct is has the capacity to interact with microtubules in a manner analogous to native tau.

We proceeded to test whether the light (488λ) induced Tau:Cry2 oligomers exhibited biochemical evidence of oligomerization using biochemical fractionation and native gel electrophoresis. The Tau::Cry2 and mCherry::Cry2 transduced cultures were exposed to light for up to 60 min. We biochemically fractionated the lysates by centrifugation, using classic approaches for isolating tau oligomers (Apicco et al., 2018; Berger et al., 2007; Santacruz et al., 2005). Immunoblots of the resulting pelleted fraction with the Tau13 antibody (recognizing all forms of tau) demonstrated a progressive increase in tau oligomers with progressive light exposure (**Figure 1E-F**), consistent with the hypothesis that light was causing the Tau::Cry2 to oligomerize.

Next, we performed non-denaturing immunoblots to evaluate whether blue light induced dimerization of the Tau:Cry2 protein. The neuronal lysates were analyzed by NativePAGE immunoblot using the human specific total tau antibody, tau13, which recognizes all tau isoforms. Increasing duration of light exposure was associated with the accumulation of high molecular weight oligomeric tau species corresponding to the expected molecular weight of dimeric oTau-c (~300 KDa); in contrast, no such accumulation of the mCherry::Cry2 was observed (**Suppl. Figure 1B, C**).

Finally, we also tested the lysates by dot blot with the anti-oligomeric tau antibodies, TOC1 (Koss et al., 2016). Increasing light exposure produced a progressive increase in TOC1 positive oligomers in the Tau::Cry2 cells (**Suppl. Fig. 1D, E**); in contrast, the companion mCherry::Cry2 cultures showed some background reactivity, but no statistically significant increases in TOC1 oligomer reactivity with progressive light exposure (**Suppl. Fig. 1D, E**). These results indicate that Tau:Cry2 oligomerizes in response to light, and that the prolonged light exposure also induces fibrillization. [Note: we use 2 antibodies to detect tau oligomers, TOC1 and TOMA2 (Combs and Kanaan, 2017; Kanaan et al., 2016; Koss et al., 2016; Ruan et al., 2020), which both recognize oligomeric tau, but are optimized for different applications: TOC1 for dot blots and immunoblots, TOMA2 for immunocytochemistry and immunohistochemistry.]

### Light induced Tau::Cry2 aggregates propagate pathology

Seeding is an important criterion for judging the presence of pathological tau. Tau sensor lines have been developed as a powerful tool for detecting the presence of tau aggregates capable of propagating pathology (Holmes et al., 2014; Sanders et al., 2014). We proceeded to test whether the light (488λ) induced Tau::Cry2 oligomers were capable of seeding. The Tau::Cry2 and mCherry::Cry2 transduced neuron cultures were exposed to light for 20 or 60 min, 10 μg lysate was applied the tau sensor line and FRET reactivity evaluated after 24 hrs; fractions from 9 month PS19 P301S tau mouse brain (S1p tau oligomer and P3 tau fibril, 10 μg each) were used as positive controls for comparison. Robust seeding was induced in the sensor line by lysate from the Tau::Cry2 60 min light exposure condition, similar to that observed upon application of tau fibrils (P3 fraction) from PS19 P301S tau mouse brains (**Fig. H, I**). Lysate from neurons expressing Tau::Cry2 exposed to 20 min light produces relatively little seeding reactivity, similar to that observed for tau oligomers isolated from PS19 P301S tau mouse brains in the S1p fraction (**Fig. H, I**). These data suggest that prolonged light exposure induces tau to pass from an oligomeric state into a fibrillar state.

Next, we directly tested for the presence of tau fibrils by staining the Tau::Cry2 cultures with thioflavine S, a histological agent that labels amyloids, including tau fibrils. Histological evaluation of the Tau::Cry2 neurons exposed to 60 min of blue light (488λ) showed the presence of inclusions positive for thioflavine S, while Cry2::Olig neurons showed no thioflavine S reactivity (**Suppl. Figure 1F, G**).

These data demonstrate that Tau::Cry2 provides the capability of rapidly inducing tau oligomerization in a manner that begins as reversible oligomers, which evolves into stable, poorly reversible aggregates of tau that exhibit thioflavine S reactivity and the ability to propagate aggregation of tau K18 fragments in the tau sensor line. The oligomers exhibit biochemical characteristics and immunochemical reactivity similar to those of native oligomers seen *in vivo*, including dimerization, reduced solubility and TOC1 reactivity (Koss et al., 2016).

### Tau oligomerization elicits phosphorylation at disease-associated epitopes

Analysis of tau phosphorylation provided insight into the relationship between oligomerization and phosphorylation in the Tau::Cry2 system. Light mediated tau oligomerization induced robust phosphorylation on S262 (**Suppl. Figure 2A, B**), in KXGS motif of the 4th repeat, detected with antibody 12E8, and T181 (**Suppl. Figure 2C-D**), in the projection domain, detected with antibody AT270); light did not increase tau phosphorylation in cells expressing just mCherry:Cry2Olig (**Suppl. Figure 2B, D**). In addition, no increase in pS202 phosphorylation was observed with the antibodies with CP13 or AT8 (data not shown), which could reflect differential signaling or steric hinderance. The kinetics of phosphorylation were markedly slower than the kinetics of oligomerization, beginning to accumulate only at 20 minutes with robust accumulation occurring only after 40 minutes of light stimulation (**Suppl. Figure 2C, D**). Notably, Tau phosphorylation accumulated both in neurons expressing Tau::Cry2 as well as in adjacent neurons, perhaps reflecting a transcellular response to the light induced tau oligomerization since it was not observed in mCherry::Cry2Olig cultures exposed to light. These data show that light-induced Tau::Cry2 tau oligomerization is sufficient to elicit tau phosphorylation.

### Light-induced Tau oligomerization elicits neurotoxicity

Tau oligomers are thought to induce neurodegeneration. We proceeded to test whether light-induced tau oligomers also are capable of inducing neurodegeneration. Cortical neuron cultures were transduced with Tau::Cry2 or mCherry::Cry2 and exposed to light (488λ) for 20 min each day for 3 days continually (**Figure 2A**). The cultured neurons were then examined using 3 independent measures of neurodegeneration: neurite length, LDH release and caspase 3 reactivity (**Figure 2B-G**).

**Figure 2.**
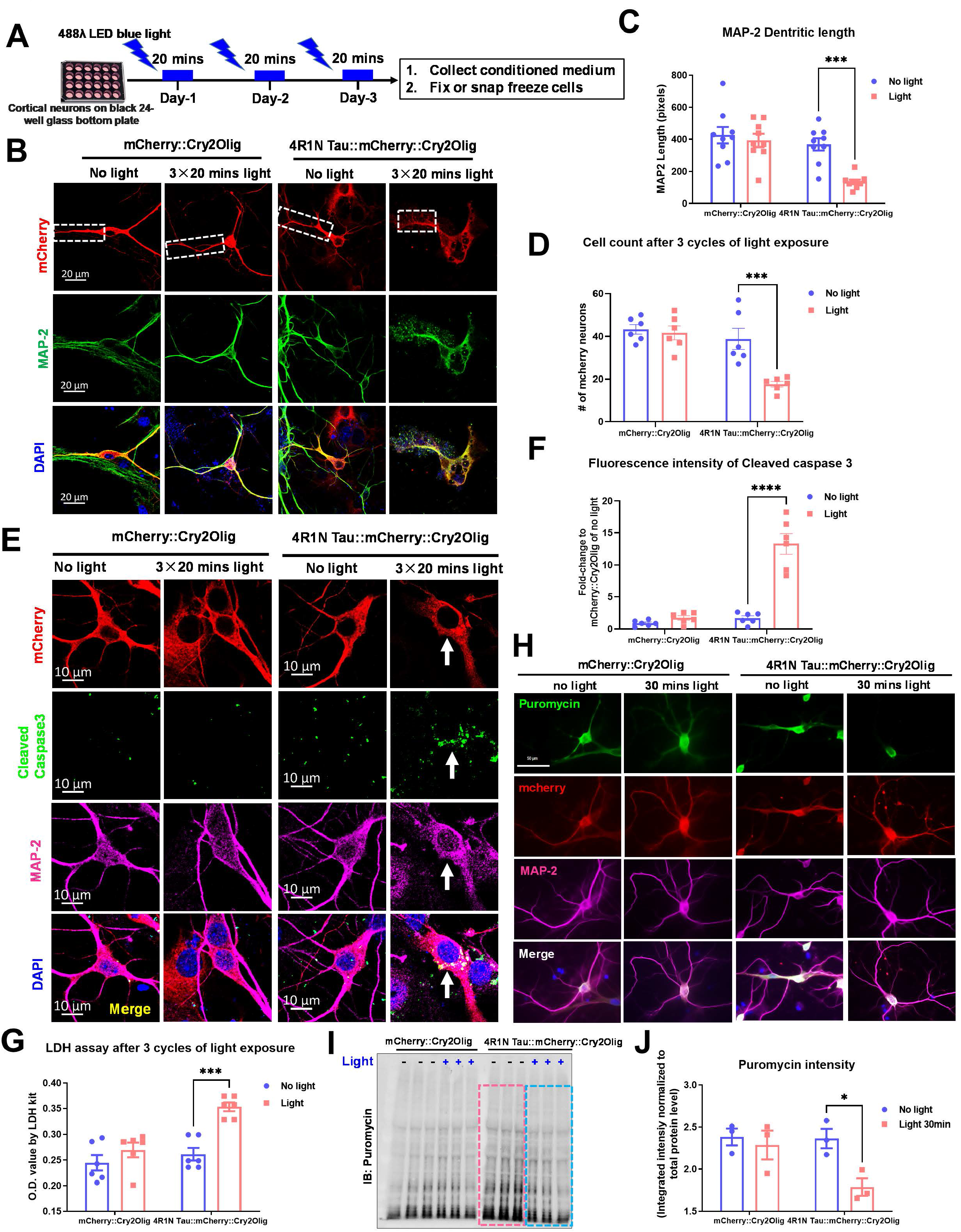
Long-lasting Tau oligomers result in neuronal toxicity. (A) Experimental diagram showed the repeated light exposure 20mins daily for continuously 3 days. This experiment was aimed to recapitulate the chronic accumulation of tau oligomers in neurons and its neurotoxicity. The cells were harvested by snap freeze or being fixed by 4% PFA according to purpose of the experiment. (B) Representative images showed the morphology of neurons over-expressed with mCherry::Cry2 or 4R1N Tau::Cry2 chimeras in the presence of long-time light exposure. Red represents mCherry positive transduced cells and green represents MAP-2 positive neurons. Scale bar, 20μm. (C) Measurement of MAP-2 positive dendritic length. The longest neurite was chosen from each neuron to measure as its dendritic length. N=8, Error bars = SEM. Two-way ANOVA with Tukey’s multiple comparisons test was used, \*\**p*<0.01 compared to control group of no light. (D) Quantification of mCherry and MAP-2 positive cell number after 3 cycles of blue light exposure. Data was from 6 independent experiments. Error bars = SEM. Two-way ANOVA with Tukey’s multiple comparisons test was performed, \*\**p*<0.01 compared to control group with no light exposure. (E) Measurement of LDH in the conditioned medium of the cell culture at 3 days after light exposure. Data collected from 6 independent experiment, expressed as O.D. value in the plate reading. Error bars = SEM. Two-way ANOVA with Tukey’s multiple comparisons test was performed, \*\**p*<0.01 compared to control group of no light. (F) Representative images showed neuronal apoptosis in 4R1N Tau::Cry2 over-expressed cells after 3 cycles of light exposure. Apoptosis was labeled by cleaved caspase 3 (green), transduced neurons was labeled by mCherry (red) and positive neurons was labeled by MAP-2 (violet). Scale bar, 10μm. (G) Quantification of the fluorescence intensity of cleaved caspase 3 in *(F)*. Data were collected from 5 neurons in each well of 5 independent experiments and normalized to the percentage of maximal fluorescence intensity. Error bars = SEM. Two-way ANOVA with Tukey’s multiple comparisons test was performed, \*\**p*<0.01 compared to no light control group. (H) Immuno-labeling of newly synthesized proteins in neurons expressing mCherry::Cry2Olig or 4R1N tau::mCherry::Cry2Olig. New proteins are labeled by puromycin (green), transduced cells labeled by mCherry (red), and neurons by MAP-2 positive (violet). Scale bar, 50μm. (I) Immunoblot of puromycin show the newly synthesized proteins during the 30 mins of incubation in cell culture with/without light exposure. (J) Quantification of puromycin intensity on immunoblots normalized to total protein levels detected by bicinchoninic acid assay. Data collected from 3 independent experiments. Error bars = SEM. Two-way ANOVA with Tukey’s multiple comparisons test, \**p*<0.05.

Neurons bearing oTau-c and exposed to light over a prolonged time course (20 min/day x 3 days) exhibited striking reductions in dendritic length (**Figure 2B, C**), aberrant dendritic morphology (**Figure 2B**) and eventually reduced cell number, which is indicative of cell death (**Figure 2D**). Immunofluorescence labeling demonstrated that the neurons exhibited increases in cleaved caspase3 intensity in light-exposed 4R1N Tau::Cry2 and MAP-2 positive neurons, indicating increased pro-apoptotic activity in the tau transduced, light exposed oTau-c expressing neurons (**Figure 2E, F**). Finally, conditioned medium collected from Tau::Cry2 (and mCherry:Cry2) transduced cultures exposed to 488λ blue light exhibited elevated levels of lactic acid dehydrogenase (LDH), which quantifies cytotoxicity (**Figure 2G**). These results provide multiple independent lines of evidence chronic induction of oTau can elicit neurodegeneration and show that oTau-c exhibits cytotoxicity in a manner reflecting the cytotoxicity of oTau, which we have shown previously (Jiang et al., 2019).

The association between oTau-c and m^6^A suggests a role in translational regulation. In order to test whether oTau-c regulates the translational stress response we monitored protein synthesis using SUnSET (Koren et al., 2019); 10 μg/ml of puromycin (a protein synthesis inhibitor that covalently incorporates into nascent proteins) was added to Tau::Cry2 or mCherry::Cry2 cortical neurons prior to light exposure, and protein synthesis was monitored with anti-puromycin antibody. Neurons transduced with Tau::Cry2 and 488λ blue light exhibited significantly decreased protein synthesis throughout the cytoplasm and neuronal arbors in comparison to the mCherry-Cry2 transduced group (**Figure 2H**). Analysis of puromycin incorporation by immunoblot normalizing to total protein levels (detected by Ponceau S) also showed a strong reduction in protein synthesis in light-exposed Tau::Cry2 neurons (**Figure 2I, J**). Thus, oTau-c inhibits protein synthesis, which is a major output of the translational stress response.

### Proteomic profiling reveals the evolution of protein interactions during tau oligomerization

We proceeded to use the power of the Tau::Cry2 system to identify the neuronal proteins that selectively bind tau oligomers. To explore the oTau protein-protein interaction (PPI) network, we performed mCherry co-immunoprecipitations (co-IP) experiments, pulling down multicomponent complexes containing the 4R1N Tau::Cry2 or mCherry::Cry2 chimeras after 0, 20 or 60 min of light exposure. The samples were then analyzed by precision Orbitrap mass spectrometry to determine the resulting PPI networks (**Figure 3A**). The label-free-quantitative protein intensities were normalized to the corresponding mCherry::Cry2 intensity in each sample (N=3/condition). The non-specific mCherry:Cry2Olig binding was subtracted from the corresponding 4R1N Tau::Cry2 for each protein in each group, and the relative protein intensities analyzed for statistical significance.

**Figure 3.**
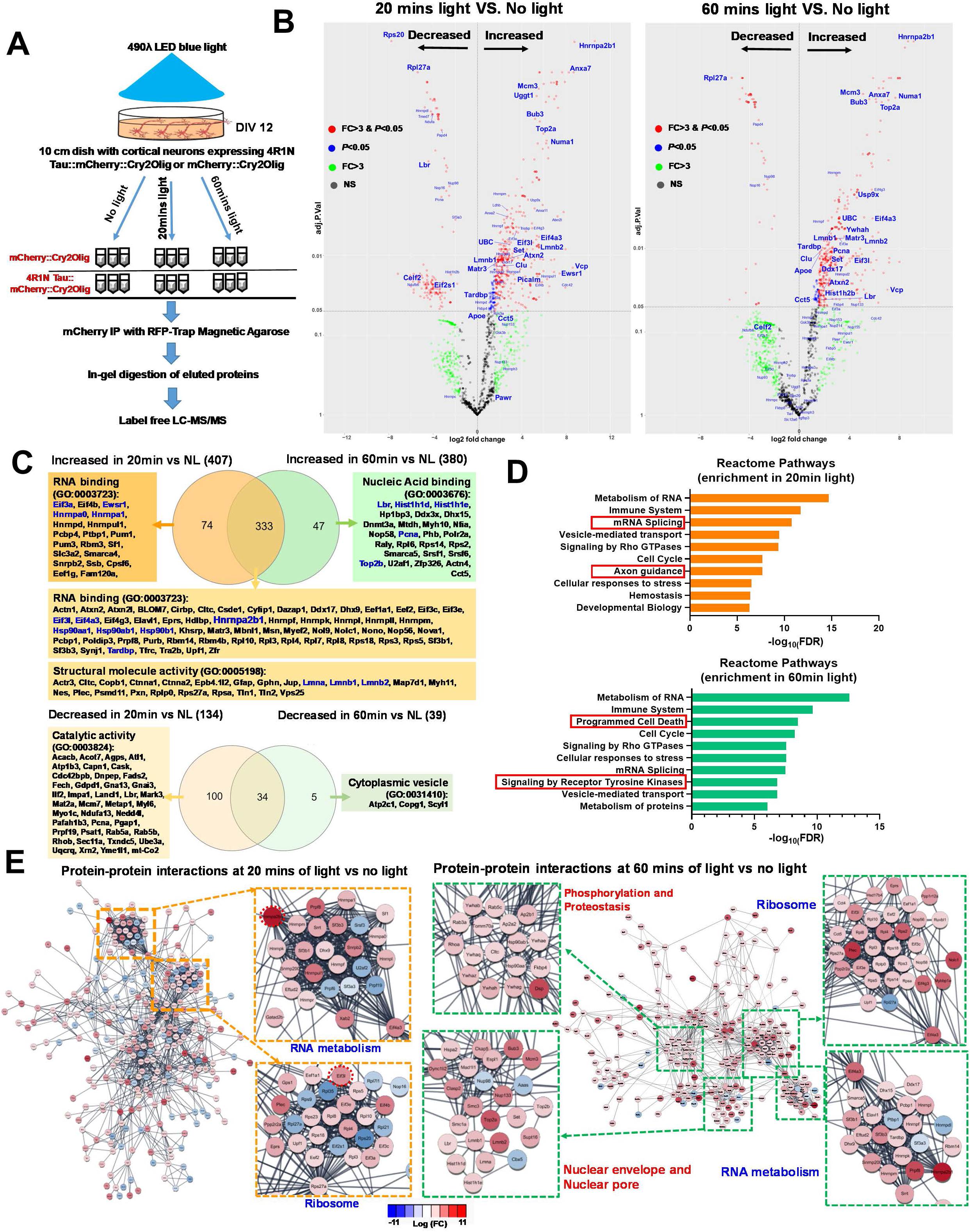
Proteomic profiling revealed the evolution of protein-protein interactions in the process of tau oligomerization. (A) Experimental design for the mass spectrometry assay to profile the protein-protein interactions at different stages of tau oligomerization. 10cm petri dishes with primary cortical neurons transduced with mCherry::Cry2 or 4R1N Tau::Cry2 lentivirus were used. These samples were snap frozen following exposure to no light or after 20mins or 60mins of 488λ blue light exposure. Cell lysates were collected and mCherry immuno-precipitation was performed with RFP-trap manganese beads. The eluted proteins were purified with in-gel digestion followed by the running of nano-LC mass spectrometry. Each condition was processed with triplicate samples. (B) Volcano plot showed the relative abundance of increased and decreased proteins that were bond to 4R1N Tau::Cry2 chimeras at 20min or 60min of light exposure in comparison to no light control group, respectively. The fold-change was calculated for individual proteins by the ratio of relative protein abundance in 20 and 60mins light to the control group of no light. Proteins significantly increased or decreased more than 3 folds (*p*≤ 0.05 and absolute log_2_ FC ≥1.58) are shown in red, whereas those exhibited significantly change but less than 3 folds (*p*≤ 0.05 but absolute log_2_ FC < 1.58) are shown in blue. Proteins showing variable changes > 3-fold were marked as green and those without significant change were marked as gray. (C) Venn diagram and enrichment of unique proteins that showed significant increased or decreased binding to 4R1N Tau::Cry2 chimeras after oligomerization triggered by 20min or 60min of blue light, respectively. Venn diagrams were drawn with increased binding proteins and decreased binding proteins respectively. The Gene ontology (GO) analysis of the unique protein in 20 min or 60min were analyzed by STRING database and the molecular function GO term and code were shown as indicated. (D) The functional analysis of proteins that exhibit significantly enriched binding to 4R1N Tau::Cry2 chimeras oligomers at early (20 mins) and late (60 mins) stages of oligomerization. Reactome pathway was analyzed from the STRING database and the top 10 Reactome pathways identified with the lowest false discovery rates (FDR) are displayed. (E) STRING network analysis of proteins that have significantly enriched binding to the complex of 4R1N Tau::Cry2 chimeras oligomers at 20min and 60mins of light exposure, respectively. Proteins were filtered by *p*<0.05 and absolute log_2_FC≥1.58 (more than 3-fold change). Only interactions with a STRING score ≥ 0.8 are shown. Evidence of interaction is represented by the distance between nodes, with more tightly packed nodes having a higher STRING score. Proteins that did not display interactions are not shown. Node color are linearly related to fold-change. At 20 mins of light, RNA metabolism and ribosomal clusters contain tightly packed nodes and are depicted magnified in the inserts. At 60mins of light exposure, in addition to RNA metabolism and ribosomal clusters, nuclear pore proteins and programed cell death signaling proteins are also clustered as shown of the magnified nods also.

Volcano plots characterizing the tau oligomer interactome network at 20 min vs. no light (**Figure 3B**, left panel), 60 min vs. no light (**Figure 3B**, right panel) and selective for 60 vs. 20 min (**Suppl. Figure 3**) identify proteins that selectively bind oTau. The Heterogeneous Nuclear Ribonucleoprotein A2/B1 (HNRNPA2B1) stands out as a major interactor with oTau-c at both 20 and 60 min, identifying a key RBP binding partner for tau; HNRNPA2B1 stands out as the major oTau-c binding protein because it is the top oTau-c binding protein at 20 and 60 min, it is the only RBP in the top 10 binders at 20 and 60 min, and mutations associated with HNRNPA2B1 dysfunction cause neurodegeneration (familial ALS) (**Supplemental Table 1**) (Kim et al., 2013a). VCP, which is another ALS-linked protein, showed up as the 3^rd^ ranked oTau-c binding protein at 20 min, which is important both because of the disease-relevance and because VCP is known to mediate tau removal, which points to a direct feedback loop for regulating oTau-c accumulation (Abisambra et al., 2013; Johnson et al., 2010; Kim et al., 2013b; McEwan et al., 2017). The nuclear envelope proteins Lamin A, B1 and B2 also bind selectively to oTau at both 20 and 60 min, but the Lamin B receptor (LBR) stands out because it shows increased binding between 20 and 60 min. Thus, oTau-c binds strongly to both lamins and LBR only at 60 min, suggesting that this interaction might play a key role contributing to the nuclear envelope disassembly that occurs during the toxic phase of tau oligomerization.

Gene ontology (GO) analysis using the STRING functional annotation database demonstrated surprising differences in the tau network components (Szklarczyk et al., 2019). A Venn diagram was created to characterize the composition of the tau binding protein networks at each time point (**Figure 3C**). Tau oligomerization at 20 min is largely reversible and shows a strong increase in the enrichment of RBPs (**Figure 3C**). Increased binding to RPBs is consistent with formation of stress granules and induction of the translational stress response, as described above. In contrast, prolonged tau oligomerization (60 min time point) is characterized by the accumulation of irreversible tau oligomers. The major increases in GO annotation groups associated with the toxic phase fall into the categories for programmed cell death and signaling by receptor tyrosine kinases (**Figure 3D**); axon guidance disappears from the top 10 categories, as might be expected in response to neuronal injury.

Enrichment analysis of the PPI network using the String database also confirmed a global shift in the oTau interaction network between 20 and 60 min. At 20 min, the most enriched categories are ribosome and RNA metabolism, while at 60 min nuclear envelope components (LBR, LMNA, LMNB1, LMNB2, Nup93 and Nup133) were prominent among the most enriched categories (**Fig 3C, E**). This shift occurs in parallel with dysfunction of the nuclear envelope and concomitant binding of multiple nuclear proteins including PCNA, POLR2A, TOP2B, HIST1H2B and HIST1H1D (**Figure 3B, E** and **Suppl. Figure 3C, D** and Suppl. Table 2), many of which function in pathways associated with lamin and LBR action (Frost et al., 2016; Maraldi, 2018). These results imply that oligomerization induces striking shifts in the tau PPI network. The delayed association with proteins linked to nuclear envelope and cell death signaling identifies novel mechanisms through which oligomeric tau induces neuronal degeneration (**Figure 3E** and **Suppl. Figure 3C, D**).

Comparing tau PPI networks among multiple studies (including our own) identifies tau binding proteins that show up most consistently (Evans et al., 2019; Maziuk et al., 2018; Wang et al., 2017; Wang et al., 2019). Variation among studies is expected because of differences in techniques, tissues, or a selective focus on a particular tau species, such as oTau. The comparisons indicate that 32 interactors were present in at least 3 of the 5 tau PPI networks, including RBPs such as EWSR1, HNRNPA’s (a0, k, r), EEFs (1a1 and 2), PCBP1 and ribosomal proteins (RPS3, 14, 27a, RPL8) (Suppl. Table 2-3). The presence of shared tau binding proteins across platforms suggests that oTau-c and oTau share similar patterns of biochemical interactions.

### HNRNPA2B1 associates with oTau and tau pathology *in vitro* and *in vivo*

The associations between proteins emerging in the proteomics dataset with oTau-c as well as native oTau were validated by immunoprecipitation (IP) and immunoblotting. Tau:mCherry:Cry2Olig was immunoprecipitated with anti-mCherry antibody from transduced cortical neurons following 0, 20 or 60 min of exposure to 488λ light. The resulting complexes were immunoblotted to detect the abundance of significant targets identified in the proteomic dataset including HNRNPA2B1, Heterogeneous Nuclear Ribonucleoprotein H (HNRNPH) and Eukaryotic Translation Initiation Factor 3 Subunit L (EIF3l) (**Figure 4A-D**). Interactions of oTau with the nuclear envelope proteins LMNB2 and LBR were also validated, as described below (**Figure 4A-D**). The proteins HNRNPA2B1 and LBR were chosen because of their strong selectivity for the toxic phase of tau oligomerization (60min) and absence of prior reports showing such interactions; while HNRNPH, EIF3l were chosen as proteins lower in the volcano plots but never-the-less exhibiting robust signals combined with strong representation at 60 min. As predicted by the proteomics, HNRNPA2B1, HNRNPH and EIF3l were strongly enriched with tau oligomerization, with respective levels being 20 min > 60 min > 0 min (**Figure 4A-D**).

**Figure 4.**
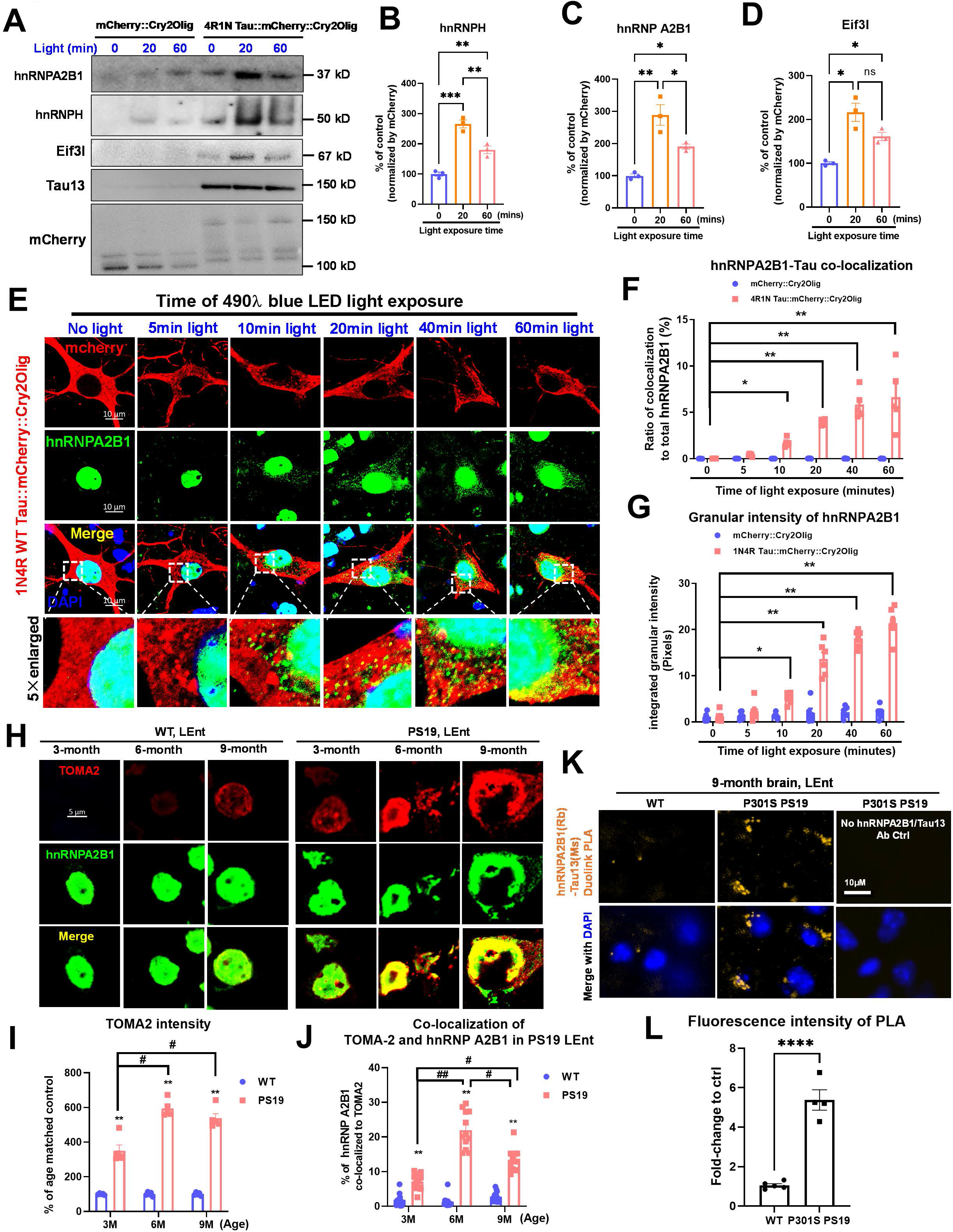
Tau oligomerization elicits striking change in HNRNPA2B1 localization. (A) Validation of the mCherry IP samples by immunoblot, including HNRNPA2B1, HNRNPH and Eif3l, respectively. Human tau antibody Tau13 was used to verify the specificity of mCherry IP in the 4R1N Tau::Cry2 over-expressed samples. Antibody to mCherry was used to detect the loading amount of mCherry::Cry2 containing cell lysate in each group as the internal control. (B) Quantification of the HNRNPA2B1 band intensity. The integrated intensity of each HNRNPA2B1 band was first normalized by the band intensity in its corresponding mCherry labeling and then was compared in its ratio to control group with no light exposure. Data were collected from 3 independent experiments. Error bars = SEM. Two-way ANOVA with Tukey’s multiple comparisons test, \*\**p*< 0.01, \**p*<0.05. (C) Quantification of the HNRNPH band intensity. The integrated intensity of each HNRNPH band was first normalized by the band intensity in its corresponding mCherry labeling and then was compared in its ratio to control group with no light exposure. Data were collected from 3 independent experiments. Error bars = SEM. Two-way ANOVA with Tukey’s multiple comparisons test, \*\**p*< 0.01, \**p*<0.05. (D) Quantification of the Eif3l band intensity. The integrated intensity of each Eif3l band was first normalized by the band intensity in its corresponding mCherry labeling and then was compared in its ratio to control group with no light exposure. Data were collected from 3 independent experiments. Error bars = SEM. Two-way ANOVA with Tukey’s multiple comparisons test, \*\**p*< 0.01, \**p*<0.05. (E) Representative images showing the translocation and co-localization of HNRNPA2B1 with tau oligomers at 0, 5, 10, 20, 40, and 60 mins of 488λ blue light exposure, respectively. Scale bar, 10μm. (F) Quantification of tau-HNRNPA2B1 co-localization by the ratio of yellow to total green. Data from 5 independent experiments. Error bars = SEM. Two-way ANOVA with Tukey’s multiple comparisons test, \**p*<0.05, \*\**p*<0.01. (G) Quantification of cytoplasmic granular HNRNPA2B1 intensity. Data from 5 independent experiments. Error bars = SEM. Two-way ANOVA with Tukey’s multiple comparisons test, \**p*<0.05, \*\**p*<0.01. (H) Representative images showing the translocation and co-localization of HNRNPA2B1 (green) with tau oligomers (labeled by TOMA2 antibody, red) in the lateral entorhinal cortex (LEnt) of PS19 P301S tau transgenic mice at age of 3-month, 6-month and 9-month, respectively. Scale bar, 5μm. (I) Quantification for the accumulation of tau oligomers in PS19 mice compared to age-matched WT control by the total fluorescence of TOMA2 labeling. Data was collected from 5 animals and normalized to the percentage of age-matched wild type mice. Error bars = SEM. Two-way ANOVA with Tukey’s multiple comparisons test, \*\**p*<0.01 compared to age-matched WT control, #*p*<0.05 in comparison of ages in PS19 mice. (J) Quantification for the co-localization of HNRNPA2B1 with tau oligomers by the co-efficiency Pearson’s R value of green (HNRNPA2B1) to red (TOMA2). Data was collected from 5 animals and normalized to the percentage of 3-month PS19 mice. Error bars = SEM. Two-way ANOVA with Tukey’s multiple comparisons test, \*\**p*<0.01 compared to young mice at 3-month old as indicated, #*p*<0.05 in comparison of 6-month and 9-month old mice. (K) Representative images of proximity ligation assay show the co-localization of HNRNPA2B1 and tau aggregates in 9-month aged PS19 mouse brain in comparison to WT control. HNRNPA2B1 was probed using a rabbit antibody and tau aggregates were probed by mouse Tau 13 antibody. The λ_ex_ 554 nm; λ_em_ 576 nm (Cyanine 3; Zeiss Filter set 20) fluorescence activity reflects the co-localization of protein molecules within 40nm. Scale bar 10μm. (L) Quantification of the PLA assay as shown in (K). The data were collected by the total orange fluorescence intensity and normalized to the fold-change of WT control group. Data are shown as mean ± SEM. \*\*\*\**p*<0.001 by two-tailed Welch’s *t*-test.

We performed IPs from PS19 P301S tau mice in order to test whether endogenous oTau exhibits a binding pattern similar to oTau-c. The results demonstrated that oTau recognized each of the validated proteins mentioned above. Antibodies to HNRNPHA2B1, HNRPH and Eif3l were used for immunoprecipitations from 3- and 6-month PS19/P301S tau mice brain tissue. The immunoprecipitates were then immunoblotted with TOC1 antibody to detect levels of oTau (**Suppl. Figure 4A-F**). oTau co-precipitated with each of the candidate binding proteins. These results demonstrate that native tau oligomers exhibit binding patterns similar to that of Tau::Cry2 oligomers, and support the hypothesis that the PPI network identified for oTau-c robustly reflects the native association network of oTau.

### Tau oligomerization elicits cytoplasmic translation of HNRNPA2B1

Having validated select protein interactions, we proceeded to examine how tau oligomerization affects localization of these cellular factors. We hypothesized that tau oligomerization would induce cytoplasmic translocation of RNA binding proteins that are associated with a SG response. To examine this, Tau::Cry2 or mCherry::Cry2 were expressed in cultured cortical mouse neurons were exposed to 488λ light for up to 60 min, fixed and labeled with anti-HNRNPA2B1 antibody (**Figure 4E-G**). Tau oligomerization elicited nuclear-cytoplasmic translocation of HNRNPA2B1 that was robust from 10 – 60 min and exhibited progressively increasing colocalization with tau during this time course (**Figure 4E-G**). We also assessed interactions with the classic SG protein markers TIA1, PABP or eIF3η (**Suppl. Figure 5**). TIA1 began translocating from the nucleus to the cytoplasm with as little as 5 min of illumination, with robust cytoplasmic translocation and colocalization with Tau::Cry2 evident at 20 min (**Suppl. Figure 5A-B**). Beginning at 20 min of light exposure we also observed coincident co-localization with PABP and eIF3η (**Suppl. Figure 5C-F**). The cytoplasmic distribution of TIA1 was diffuse at short exposure times but well defined and distinctly granular at 40 and 60 min (**Suppl. Figure 5A-B**).

The strong co-localization between HNRNPA2B1 and oTau was also apparent *in vivo*, as shown by immunohistochemistry (**Figures 4H-J, Suppl. Fig. 5G, H**) or proximity ligation assay (PLA) (**Figures K, L**). [Note that the pathology of TIA1 and many RBPs including HNRNPA2B1 are sensitive to fixation time requiring fixation of ≤ 2 hrs for robust detection (Maziuk et al., 2018).] PS19 P301S tau mice exhibit progressive enrichment of oTau (with the anti-oTau TOMA2 antibody) in the lateral entorhinal cortex (LEnt) of PS19 mice beginning at the 3-month old animals (**Figure 4H-J, Suppl. Figure 5G, H**); some oTau reactivity was also apparent in nuclei of the older C57BL/6 wild type (WT) mice (9 months), although not earlier (**Figure 4H-J**). HNRNPA2B1 showed strong cytoplasmic translocation at 6 and 9 months in PS19 mice, colocalizing strongly with oTau at 6 and 9 months (**Figure 4H-L**). Indeed, HNRNPA2B1 colocalized better with tau pathology in this model than any other RBP studied previously (Apicco et al., 2018; Maziuk et al., 2018). Use of the PLA assay emphasized the co-localization of HNRNPA2B1 with tau oligomers (TOMA2), and also highlighted the cytoplasmic location of this complex (**Figure 4K, L**).

### HNRNPA2B1 links oTau with cytoplasmic N^6^-methyladenosine in cultured neurons and P301S tau mice

The strong interaction between oTau and HNRNPA2B1 suggests a functional coupling between the two proteins. HNRNPA2B1 has been shown to act as a reader of m^6^A in the nucleus where it regulates transcription (Alarcon et al., 2015), but its function in the cytoplasm is unknown. The link between HNRNPA2B1 and m^6^A led us to hypothesize that HNRNPA2B1 might also function in the cytoplasm as a m^6^A reader. We began investigating this by exploring whether oTau co-localizes with m^6^A after light mediated tau oligomerization.

Cortical neurons were transduced with Tau::Cry2 or mCherry::Cry2 and then exposed to 0, 20 or 60 min of 488 λ light (**Suppl. Figure 6A**). The neurons were then labeled with antibodies to m^6^A and HNRNPA2B1 for analysis by imaging and immunoblot. Light exposure induced formation of cytoplasmic puncta containing oTau-c (**Suppl. Figure 6A,** top row). Increasing duration of light exposure induced a significant (~40%) increase in the association of oTau-c puncta with m^6^A puncta (**Suppl. Figure 6A, B**). These results suggest an intimate coordination between the localization of oTau-c and cytoplasmic m^6^A during stress.

We proceeded to investigate whether oTau and HNRNRPA2B1 co-localize with m^6^A in brain tissue from P301S tau mice and human AD patients. Once again, co-localization was explored using immunohistochemistry, PLA and confirmed with immunoprecipitation. Lateral entorhinal cortex (LEnt) from 3-, 6- and 9-month P301S mice, as well as the wild type (WT) littermate controls were labeled with antibodies to m^6^A and HNRNPA2B1 and examined by PLA or immunohistochemistry (**Figure 5A-C, Suppl. Fig. 6C, D**).

**Figure 5.**
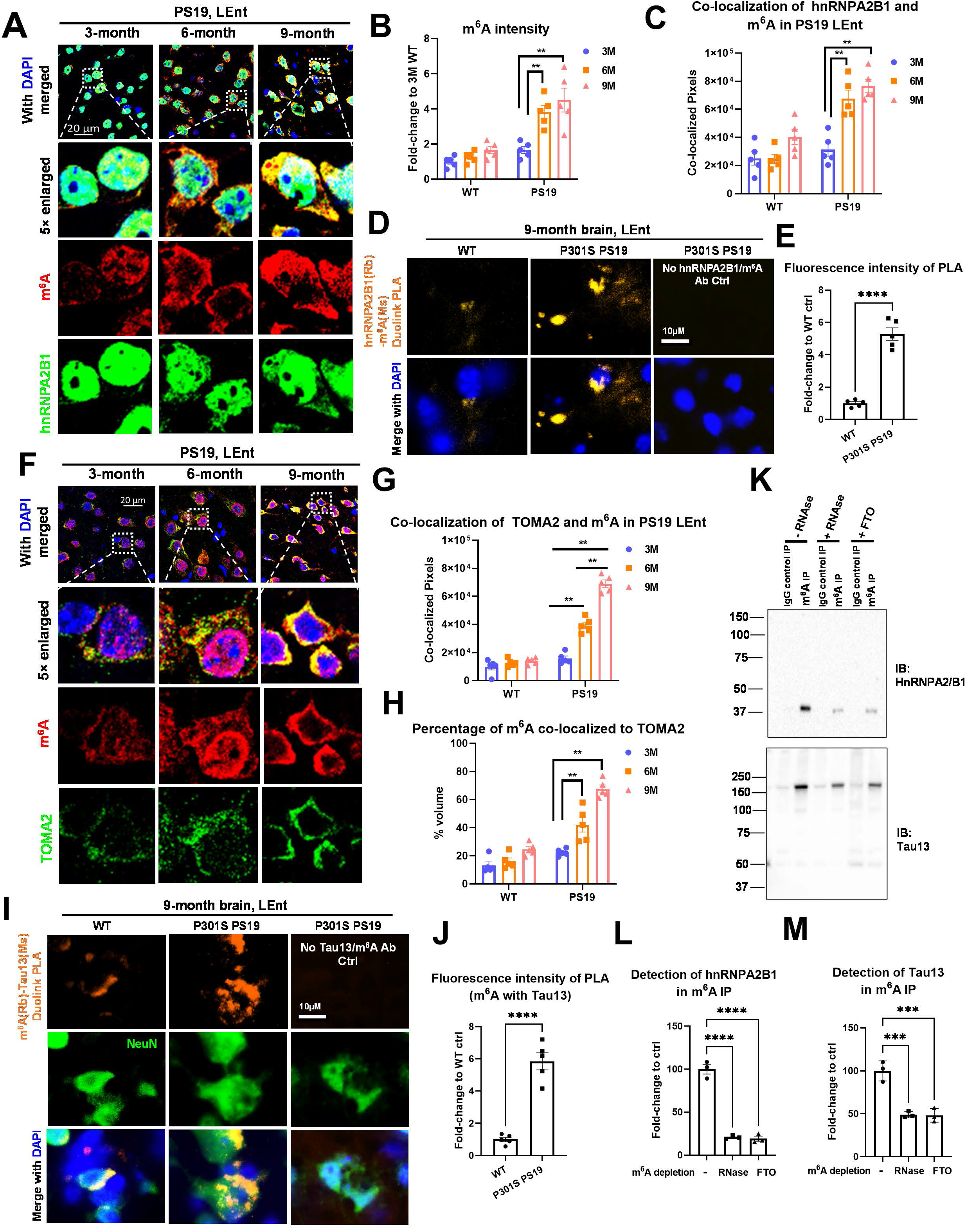
m^6^A co-localizes with HNRNPA2B1 and oligomeric tau in P301S tau transgenic mice brain. (A) Representative images showing the co-localization of HNRNPA2B1 (green) and m^6^A (red) in the lateral entorhinal cortex (LEnt) of PS19 P301S tau transgenic mice at age of 6-month and 9month, respectively. Scale bar, 20μm. (B) Quantification of the increase of m^6^A in PS19 mice compared to age-matched WT control based upon the total fluorescence of m^6^A labeling. Data was collected from 5 animals and normalized to the fold-change of 3-month wild type mice. Error bars = SEM. Two-way ANOVA with Tukey’s multiple comparisons test, \*\**p*<0.01. (C) Quantification of the co-localization of HNRNPA2B1 with m^6^A based upon the raw colocalized pixels. Data was collected from 5 animals. Error bars = SEM. Two-way ANOVA with Tukey’s multiple comparisons test, \*\**p*<0.01. (D) Representative images of proximity ligation assay (PLA) show the co-localization of HNRNPA2B1 protein with m^6^A transcripts in 9-month aged PS19 mouse brain in comparison to WT control. HNRNPA2B1 was probed by rabbit antibody and m^6^A transcripts were probed by mouse anti-m^6^A antibody. Orange fluorescence intensity reflects the co-localization of HNRNPA2B1 and m^6^A transcripts within 40nm. Scale bar 10μm. (E) Quantification of the PLA assay as shown in (D). The data was collected by the total orange fluorescence intensity and normalized to the fold-change of WT control group. Data were collected from 5 animals and is shown as mean ± SEM. \*\*\*\**p*<0.001 by two-tailed Welch’s *t*-test. (F) Representative images showing the translocation and co-localization of m^6^A (red) with oTau (labeled by TOMA2 antibody, green) in the lateral entorhinal cortex (LEnt) of PS19 P301S tau transgenic mice at age of 6-month and 9-month, respectively. Scale bar, 20μm. (G) Quantification of the co-localization of TOMA2 with m^6^A based upon the raw co-localized pixels. Data was collected from 5 animals. Error bars = SEM. Two-way ANOVA with Tukey’s multiple comparisons test, \*\**p*<0.01. (H) Quantification for the percentage of m^6^A co-localized to tau oligomers by the co-efficiency Pearson’s R value of red (m^6^A) to green (TOMA2). Data was collected from 5 animals. Error bars = SEM. Two-way ANOVA with Tukey’s multiple comparisons test, \*\**p*<0.01. (I) Representative images of PLA assay showed the co-localization of m^6^A transcripts with tau aggregates in 9-month aged PS19 mouse brain in comparison to WT control. Tau aggregates were probed by mouse antibody and m^6^A transcripts were probed by rabbit anti-m^6^A antibody. Orange fluorescence intensity reflects the co-localization of m^6^A transcripts and tau aggregates within 40nm. Scale bar 10μm. (J) Quantification of the PLA assay as shown in (D). The data were collected based on the total orange fluorescence intensity and normalized to the fold-change of WT control group. Data were collected from 5 animals and is shown as mean ± SEM. \*\*\*\**p*<0.001 by two-tailed Welch’s *t*-test. (K) Immunoprecipitation of m^6^A from 6-month PS19 brain cortex lysate. IgG antibody was used to exclude the non-specific binding. The amounts of HNRNPA2B1 and tau bound to m^6^A were detected by immunoblot. RNase or FTO group were used to deplete the m^6^A in the lysate to confirm the specific binding of HNRNPA2B1 and tau to m^6^A. (L-M) Quantification of HNRNPA2B 1 and tau band intensity as shown in (K). Data were collected from 3 independent experiments. Error bars = SEM. One-way ANOVA with Tukey’s multiple comparisons test was used, \*\*\**p*< 0.005, \*\*\*\**p*<0.001.

Immunohistochemistry showed that cytoplasmic m^6^A reactivity steadily increased in the P301S tau mice. At 3-months, occasional neurons exhibited cytoplasmic m^6^A reactivity. By 6- and 9-months, most neurons exhibited cytoplasmic m^6^A reactivity, with an intensity that was more than twice that observed at 3-months (**Figure 5A, B, Suppl. Fig. 6C, D**). The pattern of cytoplasmic HNRNPA2B1 reactivity paralleled that of m^6^A, with cytoplasmic HNRNPA2B1 appearing only in occasional neurons in 3-month P301S tau mice, and exhibiting progressively stronger cytoplasmic reactivity in 6- and 9-month P301S tau mice (**Figure 5A, B, Suppl. Fig. 6C, D**). Quantification of colocalization between HNRNPA2B1 and m^6^A followed a similar pattern, showing robust co-localization in 6- and 9-month P301S tau mice (**Figure 5A, C, Suppl. Fig. 6C, D**). For the WT mice, m^6^A reactivity was restricted to the nucleus at 3- and 6-months, while at 9-month some cytoplasmic m^6^A reactivity was apparent (**Figure 5A, B Suppl. Fig. 6C, D**). Similarly, HNRNPA2B1 reactivity was nuclear at 3- and 6-months, but exhibited a small amount of cytoplasmic reactivity in 9-month WT mice.

PLA analysis demonstrated robust reactivity for m^6^A and HNRNPA2B1 interactions in the 9-month old P301S tau mice, but only very little reactivity in 9-month old WT mice (**Figure 5D**). Notably, the m^6^A and HNRNPA2B1 PLA reactivity occurred only in the cytoplasm. These data demonstrate that m^6^A and HNRNPA2B1 exhibit strong interaction in the cytoplasm of elderly P301S tau mice.

The response of oTau paralleled the observations for m^6^A and HNRNPA2B1. A small amount of oTau reactivity (detected with the TOMA2 antibody) was apparent in 9-month WT mice and 3-month P301S tau mice, while 6- and 9-month P301S tau mice exhibited progressively more oTau reactivity. Importantly, strong co-localization was observed between m^6^A and oTau in 6- and 9-month old P301S tau mice (**Figure 5E-G**). PLA analysis again demonstrated robust reactivity for m^6^A and TOMA2 interactions in the 9-month old P301S tau mice, but only very little reactivity in 9-month old WT mice (**Figure 5E-G**).

### HNRNPA2B1 links oTau with cytoplasmic N^6^-methyladenosine in human AD cases

We next examined co-accumulation in human temporal cortex tissues (Brodmann 41/42) from age-matched AD and Normal Control cases (7 AD/7 Ctl). As expected, abundant cytoplasmic m^6^A and oTau reactivity were detected in the cytoplasm of neurons in the AD cases (**Figure 6A-C**), while little cytoplasmic reactivity was observed in the control cases (**Figure 6A-C**). Quantification of the corresponding immunofluorescence showed an increase in labeling of almost 4-fold (**Figure 6D, E**, p<0.01). The cytoplasmic oTau showed strong co-localization with the cytoplasmic m^6^A reactivity, as seen in the mouse tauopathy model (**Figure 6A-C**).

**Figure 6.**
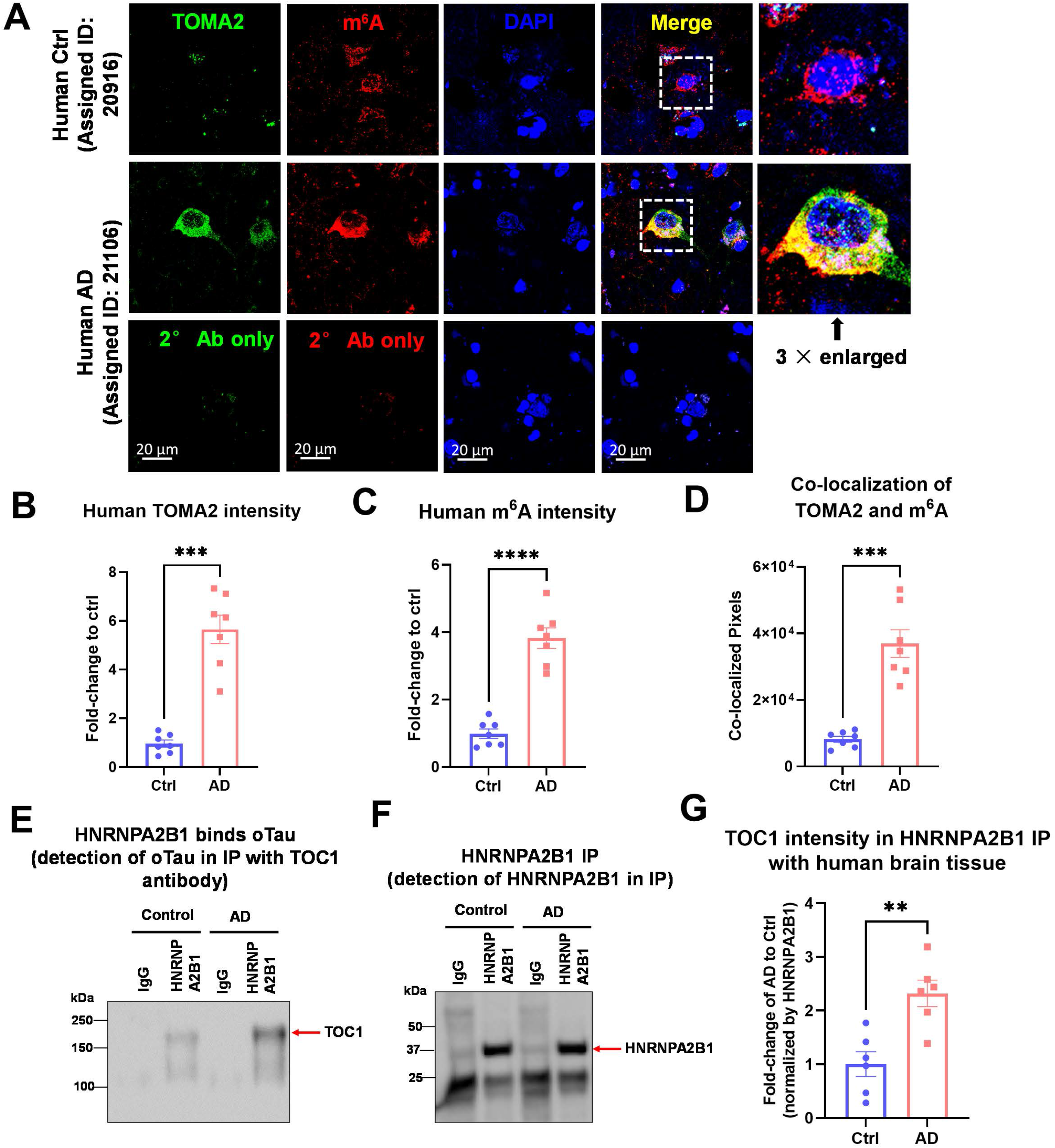
m^6^A co-localizes with oligomeric tau in post-mortem human AD brain tissue. (A) Representative images showing the accumulation and co-localization of m^6^A (red) with tau oligomers (green, labeled with the TOMA2 antibody) in post-mortem human AD brain temporal tissue in comparison to age-matched normal controls. Scale bar, 20μm. (B) Quantification of oTau accumulation in post-mortem human AD brain temporal tissue based the total fluorescence intensity of TOMA2 labeling. Data was collected from 7 human age-matched control and 7 AD cases. Data was normalized to the fold-change of control and is shown as mean ± SEM. \*\**p*<0.01 by two-tailed Welch’s *t*-test. (C) Quantification for the increase of m^6^A in post-mortem human AD brain temporal tissue by the total fluorescence intensity of labeling. Data was collected from 7 human age-matched control and 7 AD cases. Data was normalized to the fold-change of control and shown as mean ± SEM. \*\**p*<0.01 by two-tailed Welch’s *t*-test. (D) Quantification for the co-localization of TOMA2 to m^6^A shown by the raw co-localized pixels. Data was collected from 7 human age-matched control and 7 AD cases. Error bars = SEM. \*\**p*<0.01 by two-tailed Welch’s *t*-test. (E-F) Immunoprecipitation of HNRNPA2B1 from post-mortem human AD brain tissue or age-matched control. IgG antibody was used to exclude the non-specific binding. The amount of oligomeric tau bond to HNRNPA2B1 were detected using TOC1 antibody by immunoblot (E). The amount of pull-down HNRNPA2B1 was also probed by HNRNPA2B1antibody (F). (G) Quantification of oligomeric tau bond to HNRNPA2B1. Band intensity of TOC1 was normalized to the HNRNPA2B1 band. Data was collected from 7 human age-matched control and 7 AD cases. Error bars = SEM. \*\**p*<0.01 by two-tailed Welch’s *t*-test.

We proceeded to test for biochemical evidence of interaction between m^6^A, HNRNPA2B1 and tau oligomers in human frontal cortex tissues (Brodmann 10) using age-matched AD and Normal Control cases (6 AD/6 Ctl). oTau in HNRNPA2B1 immunoprecipitates were observed to be increased over 2-fold in AD compared to control brain (**Suppl. Figure 7A-C**), despite pulling down similar amounts of HNRNPA2B1 from AD and Ctrl tissues (**Suppl. Figure 7B**). Thus, AD brain exhibits increased formation of stable complexes containing oTau and HNRNPA2B1.

### HNRNPA2B1 is required for the association of tau oligomers with m^6^A

We used a genetic knockdown approach to explore whether HNRNPA2B1 was important for the interactions between oTau and m^6^A in the cytoplasm. Neurons expressing Tau::Cry2 or mCherry::Cry2 were subjected to knockdown with siRNA coding for HNRNPA2B1 or control. Knockdown of HNRPA2B1 was approximately 50% (**Figure 7A-D**). Because m^6^A and oTau both associate with SGs, we examined whether HNRNPA2B1 knockdown impacted on the association between oTau-c and m^6^A. Imaging m^6^A showed that light induced tau oligomerization increased the amount of oTau-c associated cytoplasmic m^6^A (**Figure 7A**, green channel). We compared the results with knockdown of HNRNPA2B1 and observed a marked reduction of the association of oTau-c with the m^6^A puncta compared to control knockdown, decreasing the association between oTau and m^6^A puncta by about 40% (**Figure 7A, D**). We hypothesized that the changes in association of oTau-c with m^6^A might be reflected by changes in translation, because both oTau and m^6^A are known to regulate RNA translation (Anders et al., 2018; Chen et al., 2019).

**Figure 7.**
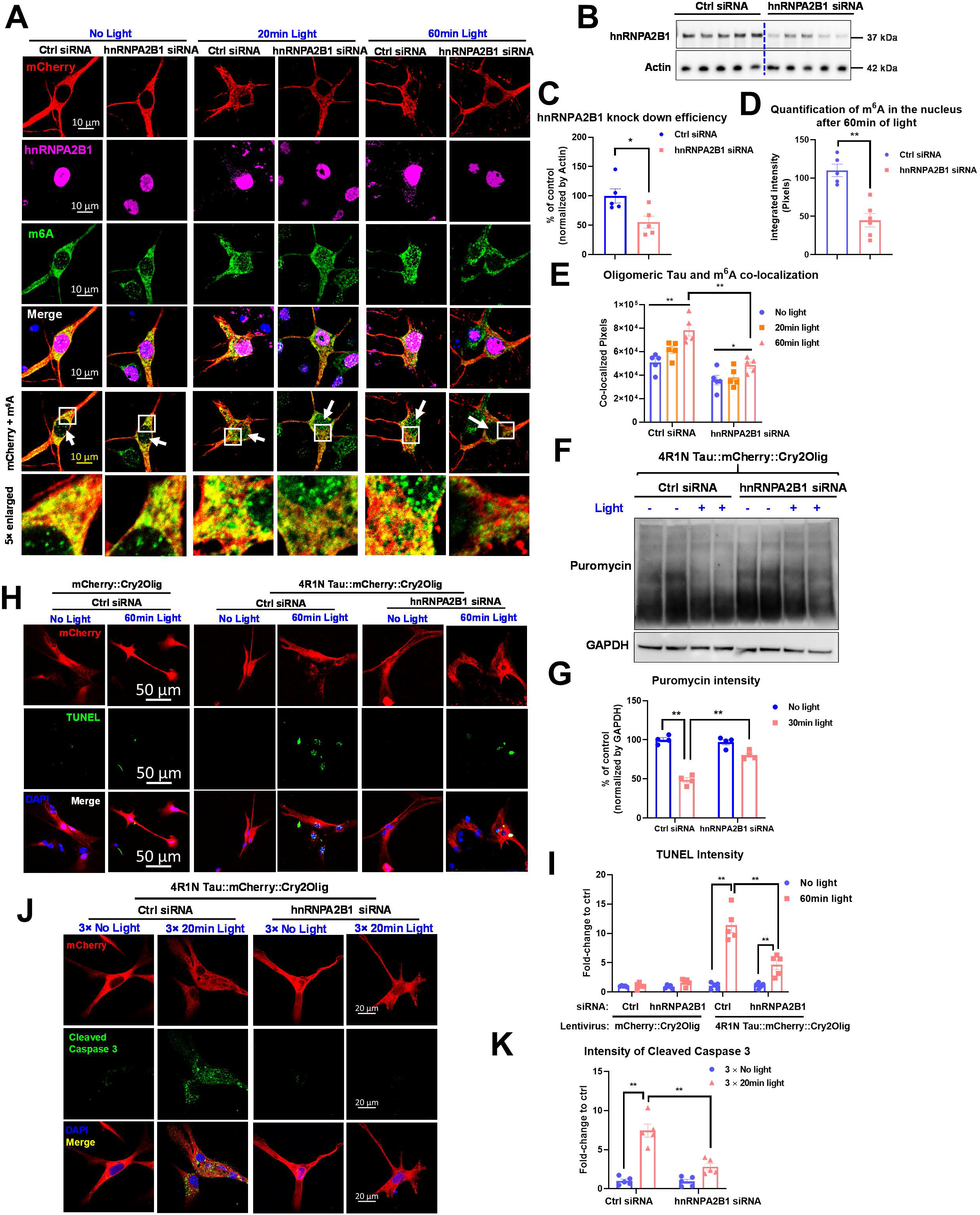
HNRNPA2B1 knockdown reduces oTau-c induced translational stress, DNA damage and association with N^6^-methyladenosine RNA. (A) Images showing the effects of HNRNPA2B1 knockdown on the association between oTau-c and m^6^A. Neurons were transduced with Tau::Cry2 (red) plus siRNA directed against HNRNPA2B1 or scrambled control. The neurons were then exposed to 488 nm light for 0, 20 or 60 min, fixed and labeled for HNRNPA2B1 (purple), m^6^A (green) or DAPI (blue). Scale bar 10 μm. (B, C) Quantification of siRNA mediated knockdown of HNRNPA2B1. B: Immunoblot of lysates run on a 12% reducing gel and probed with anti-HNRNPA2B1 antibody and actin antibody as the internal control. C: Quantification of band intensities from the immunoblot demonstrates a 45% reduction of HNRNPA2B1protein in the primary cortical neuron cultures. \**p*<0.05 by a two-tail *t*-test, N=5, mean ± SEM. (D) Quantification of the colocalization between oTau-c and m^6^A. Image analysis shows that HNRNPA2B1 knockdown elicited a strong reduction in the fraction of oTau-c colocalized with m^6^A, which is apparent in the reduction of yellow pixels evident visually or by scatterplot (bottom row). HNRNPA2B1 knockdown produced a statistically significant reduction in oTau-c - m^6^A colocalization at every time point when analyzed quantitatively, Two-way ANOVA with Tukey’s multiple comparisons test, \**p*<0.05, \*\**p*<0.01. (E) Quantitative imaging demonstrates that HNRNPA2B1 knockdown produced a statistically significant reduction in nuclear m^6^A puncta after 60 min of light exposure. Unpaired T-test with Welch’s correction, two-tailed, \*\**p*<0.01. (F) Immunoblot of puromycin showing the newly synthesized proteins in conditions with siRNA towards HNRNPA2B1 or scrambled control in conditions with no light or 30min light. GAPDH antibody was probed as the internal control. (G) Quantification of puromycin band intensities, which were internalized by GAPDH band intensity before being normalized to the control group without light or HNRNPA2B1 knockdown. Data was from 4 independent experiments. Error bars = SEM. Two-way ANOVA with Tukey’s multiple comparisons test was used, \*\**p*<0.01. (H) Representative images of TUNEL labeling show that 60mins of continuous tau oligomerization induced DNA damage which can be ameliorated by HNRNPA2B1 knockdown. Red shows the mCherry::Cry2Olig or 4R1N Tau::mCherry::Cry2Olig positively transfected neurons. Green represents the TUNEL signaling in the nucleus and DAPI was used for the labeling of nucleus. Scale bar 50μm. (I) Quantification of TUNEL labeling intensity. Treatment with HNRNPA2B1 siRNA reduces TUNEL intensity by more than 50% (compared to siRNA control treatment) following 60 min light exposure of the Tau::Cry2 cells. Data are shown as mean ± SEM. Two-way ANOVA with Tukey’s multiple comparisons test, \*\**p*<0.01. (J) Representative images of cleaved caspase3 labeling show that HNRNPA2B1 knockdown abrogates toxicity induced by extended light exposure in Tau::Cry2 neurons. Red shows the mCherry::Cry2Olig or 4R1N Tau::mCherry::Cry2Olig positively transfected neurons. Green represents cleaved caspase 3 reactivity identifying apoptotic activity, and DAPI was used for the labeling of nucleus. Scale bar 20μm. (K) Quantification of cleaved caspase 3 labeling intensity. Data was normalized to the fold-change to their compared control group. After 3 cycles of tau oligomerization, 20 min daily, the cleaved caspase 3 intensity was increased over 6-fold while it was reduced more than 3-fold (to <33%) with HNRNPA2B1 knock down. Data shown as mean ± SEM. Two-way ANOVA with Tukey’s multiple comparisons test, \*\**p*<0.01.

Tau oligomerization also elicited strong changes in nuclear m^6^A, although no Tau:Cry2 was observed in the nucleus (**Figure 7A, E**). Tau oligomerization greatly increased formation of m^6^A puncta in the nucleus (**Figure 7A**). These results are consistent with known associations between m^6^A and nuclear transcription and chromatin structure (Alarcon et al., 2015; Chen et al., 2019; Liu et al., 2020). Such changes might be expected since tau pathology is known to induce changes in DNA function and structure (Cornelison et al., 2019; Frost et al., 2016; Sun et al., 2018). Surprisingly, knockdown of HNRNPA2B1 elicited a dramatic loss of nuclear m^6^A reactivity, suggesting a link between HNRPA2B1 and the nuclear stress response (**Figure 7A, E**).

Next, because HNRNPA2B1 is a putative m^6^A reader (Anders et al., 2018; Arguello et al., 2017; Chen et al., 2019), we tested whether HNRNPA2B1 is required for the regulation of translation by oTau-c. Neurons expressing Tau:Cry2 and siRNA for HNRNPA2B1 or control were exposed to light for 20 min, in the presence of puromycin. The resulting newly synthesized proteins were immunoblotted and quantified to measure protein synthesis (**Figure 7F, G**). Knockdown of HNRNPA2B1 greatly reduced the effects of oTau on translation (**Figure 7F, G**), demonstrating the requirement of HNRNPA2B1 in translational inhibition mediated by oTau-c.

### HNRNPA2B1 is required for tau-mediated neurodegeneration

The requirement for HRNPA2B1 for the actions of oTau-c on translation and the stress response led us to hypothesize that HNRNPA2B1 might also participate in toxicity caused by oTau-c. To test this, we examined the effects of HNRNPA2B1 knockdown on neurodegeneration occurring after persistent oTau-c accumulation. We scored for DNA damage first because the increases in nuclear m^6^A in light treated Tau:Cry2 neurons suggest changes in chromatin structure; in addition, DNA cleavage is a well-documented readout of apoptosis. Caspase activity was measured in the siRNA (HNRNPA2B1 or scrambled control) transduced cultures with antibody to cleaved caspase 3 (using immunocytochemistry and dot blot). Neurons expressing Tau:Cry2 were transduced with siRNA for HNRNPA2B1 or control, exposed to 488 light for 60 min, fixed and analyzed by the Terminal deoxynucleotidyl transferase dUTP nick end labeling (TUNEL) assay. Light only increased TUNEL reactivity in neurons expressing Tau:Cry2 (**Fig 7. H, I**). Importantly, HNRNPA2B1 knockdown greatly reduced the amount of TUNEL reactivity by >50%, indicative of neuroprotection. We also tested the effects of HNRNPA2B1 knockdown in an assay reflecting chronic conditions. Neurons expressing Tau:Cry2 treated with siRNA for HNRNPA2B1 or control were treated with light for 20 min per day for 3 days. At the end of the treatment period, the cells were fixed and labeled with antibody detecting cleaved caspase 3, which is a classic marker of apoptosis. Knocking down HNRNPA2B1 greatly reduced the amount of cleaved caspase 3 reactivity in the light exposed Tau:Cry2 neurons by >60% (**Figure 7J, K**). These results demonstrate a strong role for HNRNPA2B1 in the neurotoxicity elicited by oTau-c.

## DISCUSSION

Stress leads to tau phosphorylation and oligomerization (Silva et al., 2019; Vanderweyde et al., 2016). Persistent stress leads to the accumulation of phosphorylated, oligomeric and fibrillar tau, such as occurs in the context of neurodegenerative tauopathies (Wolozin and Ivanov, 2019). Accumulating research suggests that these post-translational changes in tau act as part of the integrated stress response, and are associated with the accumulation of stress granules and the inhibition of protein synthesis (Koren et al., 2020; Wolozin and Ivanov, 2019). The results presented above shed light on these actions of tau, and demonstrate that oligomerization of tau allows it to form a trimeric complex that links it to mRNA bearing N^6^-methyladenosine, a stress associated modification, through binding of HNRNPA2B1, which acts as a reader of m^6^A. Our studies further reveal a striking, ~4-fold increase in m^6^A levels in the AD brain. The discovery that m^6^A is increased in AD potentially reveals a previously unrecognized domain of molecular changes occurring in the Alzheimer brain.

The cytoplasmic localization of the oTau-HNRNPA2B1-m^6^A complex (TH-m^6^A complex) presents a striking contrast to prior studies, which largely focused on interactions between HNRNPA2B1 and m^6^A in the nucleus, where it regulates chromatin state and transcription (Anders et al., 2018; Arguello et al., 2017; Chen et al., 2019). These studies suggest that HNRNPA2B1 reads m^6^A when binding to mRNA is increased by m^6^A modifications that indirectly facilitate HNRNPA2B1 binding upon changing the secondary mRNA structure. However, stress causes the cytoplasmic translocation is HNRNPA2B1, which then associates m^6^A modified RNA in stress granules. Our prior work demonstrates that oTau also accumulates in SGs and facilitates formation of TIA1-positive SGs (Apicco et al., 2018; Jiang et al., 2019; Vanderweyde et al., 2016). Thus, the SG acts as a membraneless organelle that brings together these three components to form the TH-m^6^A complex.

Our studies begin to elucidate some of the functions of the TH-m^6^A complex. One function is to regulate protein synthesis. The accumulation of oTau inhibits protein synthesis, which likely occurs as part of the translational stress response. We show that silencing HNRNPA2B1 abrogates this response, allowing maintenance of protein synthesis in the face of stress. Tau and HNRNPA2B1 are both linked to neurodegeneration, and silencing HNRNPA2B1 prevents the toxicity associated with the accumulation of oTau (Hutton et al., 1998; Kim et al., 2013a; Qi et al., 2017). These data support the hypothesis that interaction of oTau with HNRNPA2B1 contributes to mechanisms of toxicity associated with oTau. It is worth noting that HNRNPA2B1 is probably not required for all the actions of oTau because it is not present in all oTau granules, nor is it present in all TIA1 positive SGs. However, the ability of HNRNPA2B1 knockdown to significantly inhibit the actions of oTau towards protein synthesis and toxicity indicate that HNRNPA2B1 is an important part of the oTau mechanistic pathway.

The interaction of oTau with HNRNPA2B1 is consistent with accumulating proteomic studies of neurodegenerative diseases in which HNRNPs appear prominently, as well as a recent study showing increases in m^6^A in the brains of AD cases (Han et al., 2020; Johnson et al., 2020; Johnson et al., 2018; Umoh et al., 2018). These multiple intersecting lines of evidence support the hypothesis that HNRNPA2B1 contributes to the pathophysiology of tauopathies by enabling oTau to control the utilization of m^6^A modified transcripts.

Use of the genetically modified oTau-c to model the actions of oTau presents a potential pitfall for our study. However, a surprising proportion of the actions of oTau-c are also observed in studies of oTau. Throughout the study we repeatedly investigated whether particular characteristics of oTau-c were also observed with native oTau. For instance, prior studies show that the filaments produced by tau constructs tagged with fluorescent proteins are thicker than native tau filaments, so we anticipate that the ultrastructure of oTau-c might differ from that of oTau (Kaniyappan et al., 2020). However, many other biochemical characteristics of oTau-c share features in common with oTau. Both oTau and oTau-c are recognized with antibodies (TOC1 and TOMA2) directed against native oTau, which suggests that oTau and oTau-c share similar conformational epitopes. As with oTau, prolonged oligomerization causes oTau-c to progress from a reversible to an irreversible oligomer that shows reduced solubility, much like oTau. oTau is well documented to cause neurotoxicity (Jiang et al., 2019; Lasagna-Reeves et al., 2011), and oTau-c also induce neurotoxicity.

The presence of many binding partners shared among total tau and oTau-c PPI networks indicates that oTau-c shares binding characteristics with native tau. The network we created identified additional cellular binding partners, each of which were also shown to interact with native oTau. Immunohistochemical, PLA and IP studies all demonstrated that these binding interactions were also observed with oTau using P301S tau mice and in human brain. The results obtained using human clinical samples enables assessment using tissues with endogenous levels of tau, which obviates potential artifacts derived from protein over-expression, and further demonstrate that the interactions between oTau, HNRNPA2B1 and m^6^A are robust pathological features of AD, and potentially other neurodegenerative disorders. All of these independent lines of evidence coalesce to support the hypothesis that the optogenetic induction of tau oligomerization faithfully models the interactions of native oTau with many other proteins. Because of the similar actions of oTau-c and oTau, the term “oTau” is used interchangeably with oTau-c throughout this discussion.

The process of oligomerization is central to cell biology and is known to regulate the functions of many proteins. For instance, oligomerization plays a critical role in the activity of p62/SQSTM, which is linked to protein aggregation in neurodegenerative diseases (Bjorkoy et al., 2005; Fecto et al., 2011; Niu et al., 2018). Monomeric p62 directs ubiquitinated proteins to the proteasome, while oligomeric p62 directs ubiquitinated proteins to the autophagosome (Wurzer et al., 2015). Oligomerization also activates the functions of p53 and other transcription factors (Fischer et al., 2016; Nguyen et al., 2019; Sun et al., 2013). The current study extends the biology of oligomerization to that of tau, pointing to a fundamental mechanism in disease progression.

In conclusion, our work suggests that tau oligomerization occurs as part of a normal physiological response to stress. oTau tethers to m^6^A modified mRNA through binding to HNRNPA2B1, a reader of m^6^A modified transcripts. The TH-m^6^A complex regulates the RNA translational stress response and mediates toxicity of oTau. Our study reveals that m^6^A and the TH-m^6^A complex in particular are increased in AD. The study of RNA modifications has opened novel roads for diagnostics and personalized medicine in cancer (Lan et al., 2019; Liu et al., 2019). Our discovery of the prominent m^6^A changes in AD, and the strong relationship to tau biology opens up similar avenues for investigation in other neurodegenerative diseases.

## METHODS

### Animals

Use of all animals was approved by the Boston University Institutional and Animal Care and Use Committee. All animals were housed in IACUC-approved vivariums at Boston University School of Medicine. Timed pregnant C57BL/6 were purchased from Charles River laboratories and delivered at E-14. The tau knockout B6.129X1-*Mapt*^tm1Hnd^/J mice were purchased from the Jackson laboratory (Stock No:007251) and breeding in-house for the postnatal day-0 pups.

PS19 P301S tau transgenic mice: PS19 mice overexpressing human P301S Tau (B6;C3-Tg(Prnp-MAPT*P301S)PS19Vle/J, stock #008169) were purchased from Jackson Laboratories. Male and female PS19 P301S tau^+/-^ mice were used as breeding pairs and the F1 generation of P301S tau^+/-^(PS19) and P301S tau^-/-^(wild type) were used for the experiment. Littermates of the same sex were randomly assigned to experimental groups. Mice were sacrificed for experiment at the age of 3, 6 and 9 months old, respectively.

The fixation method of the brain tissue is critical for the successful immuno-labeling of tauopathy and RNA binding proteins in this study. Mice were anaesthetized with Ketamine/xylazine cocktail (Contains: 87.5 mg/kg Ketamine and 12.5 mg/kg Xylazine) at 0.1ml/20g.bw by i.p. injection. The mice were then perfused through heart with 20ml ice cold PBS at the speed of 4ml/min for 5 mins, followed by perfusion with 20ml ice cold 4% PFA for 10 mins until the mouse tail became curved and stiff (note to change the speed set-up as 2ml/min when running with PFA). The mouse brains were dissected and placed in 4% PFA on ice for 1-2 hrs before it was transferred into PBS and stored at 4°C. To prepare for collecting brain sections, the fixed mice brains were transferred into 30% sucrose/PBS until the brains sank to the bottom of the tube (about 48h), and then sectioned. The fixed brains were sliced into 30μm coronal sections by cryostat, and stored in 0.005% sodium azide/PBS solution at 4°C for up to 3 months. For longterm storage, the sections were transferred into cryoprotectant solution (30% glycerol and 30% ethelyne glycol in PBS), and stored at −20 °C.

### Human brain samples

Human temporal superior gyrus tissues (Broadman areas 41/42) were used for the immunohistochemical studies. The samples were de-identified, and described in the table below. The tissue was fixed in periodate-lysine-paraformaldehyde (PLP) fixative for 2 hours, followed by overnight incubation in 30% sucrose, after which tissue sections were cut at thickness of 30 μm.

### Human cases (tissue were fixed in periodate-lysine-paraformaldehyde)

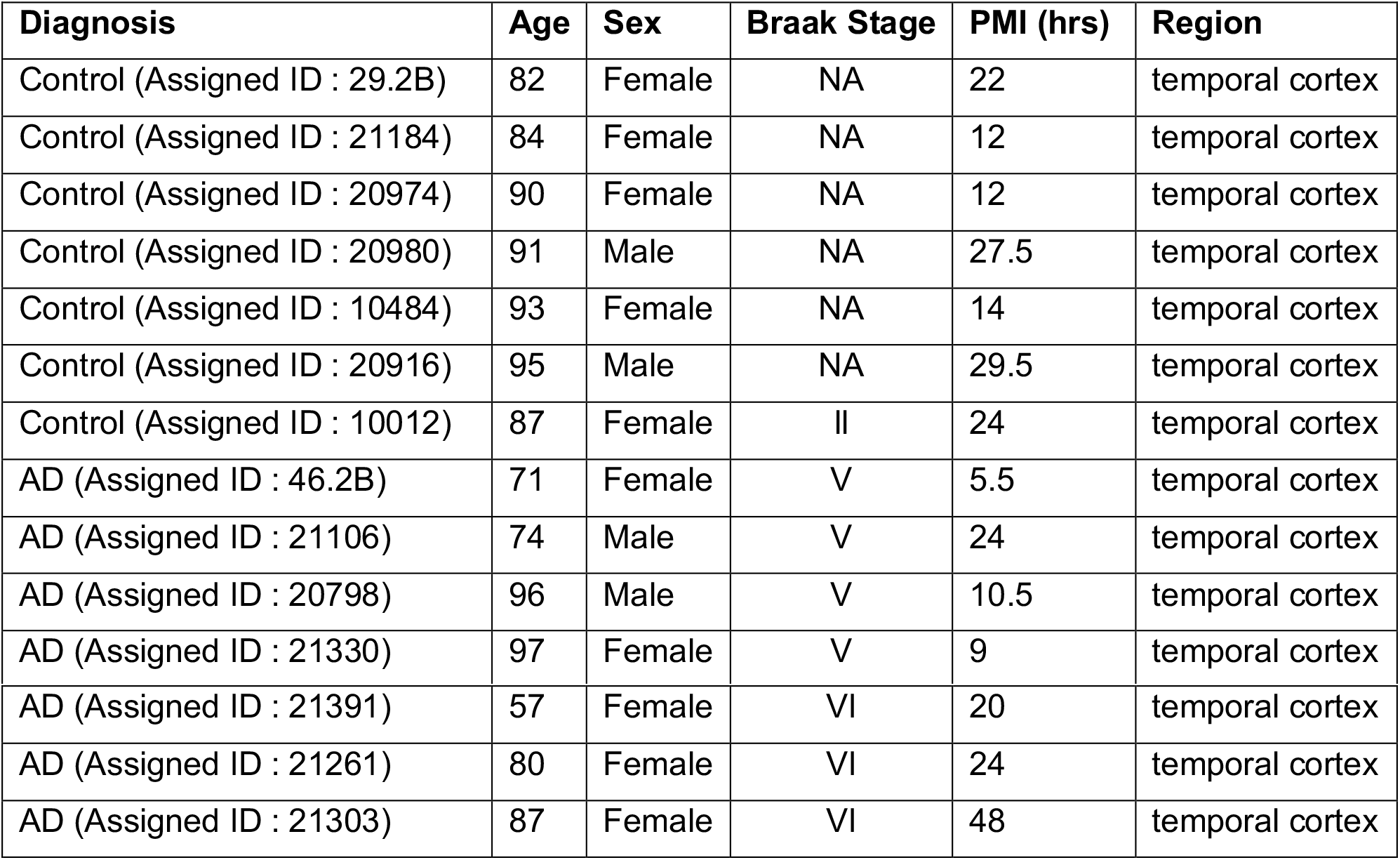

### Reagents

Prolong Gold Antifade Reagent was obtained from Thermo Fisher Scientific (Cat#P36930). NuPAGE™ 3-8% Tris-Acetate Protein Gels was purchased from Invitrogen (Cat# EA03785BOX). Novex™ HiMark™ Pre-stained Protein Standard was purchased from Invitrogen (Cat# LC5699).

### Plasmid construction

The mCherry::Cry2 and 4R1N Tau::Cry2 construct were designed by Dr. Brangwynne group. Briefly, sequences for mCherry and Cry2Olig (Addgene) were cloned into the pHR lentiviral backbone to generate the pHR-mCh-Cry2Olig plasmid. DNA fragment encoding human 4R1N full length tau were amplified by PCR using tau cDNA (Addgene). And then the DNA fragment encoding 4R1N tau were inserted into the linearized pHR-mCh-Cry2Olig backbone using In-Fusion Cloning Kit (Takara). The resulting constructs were fully sequenced to confirm the absence of unwanted substitutions.

### Lentivirus production

HEK293T cells were plated at a concentration of 1×10^6^ cells/well in a 6-well plate. 18 hrs later, cells were transiently co-transfected with PSP (1200 ng), VSV-G (400 ng), and target (400 ng) plasmids using 6 μL FuGene HD (Promega, Cat# E2311). 72 hrs later, conditioned media was harvested and centrifuged at 1000xg for 5 min to remove dead cells and debris. Supernatant was stored at −80°C until use. For primary neuron transduction, lentivirus was concentrated 10x using lenti-X concentrator (Takara Bio USA, Cat# 631231) with the concentrated pellet being resuspended in PBS with 25 mM HEPES, pH 7.4.

### Primary cortical culture with P0 pups

Primary cortical cultures were generated from postnatal P0 pups. For culturing primary cortical neurons, fresh cortical tissues were dissected from postnatal day-0 pups. Briefly, C57BL/6 pups were anesthetized via hypothermia by wrapping in gauze and placing in aluminum foil pouch on ice. Then the pups were sprayed with 70% ethanol and transferred to 60 cm dish. We then isolated the brain from the skull and separated the two hemispheres of cortex from the midbrain and transferred them to 10 cm culture dish filled with HBSS dissection buffer. The meninges of the cortex tissue were them completely removed. We them transferred all the cortical tissue into a 15 mL conical tubes and replaced HBSS dissection buffer with 5ml 0.25% Trypsin-EDTA supplemented with 150 μL DNase. And then the tissue was incubated in 37°C water bath for 15 min. And then the tissue was then carefully removed of trypsin and gently washed 3 times with HBSS dissection buffer followed by spinning down in 2000rcf for 2 mins room temperature. The tissues were resuspended in 25ml plating medium (MEM Gibco #11090, 2.5% FBS, 1x Penicillin/streptomycin, L-glutamine, 0.6% D-glucose) and triturate gently with 5 ml pipette. Single cells were passed through a 70 μm cell strainer and cell number counted. Based on the purpose of the experiment, the cortical cells were then plated on different types of culture plate or dishes with feeding medium (Neurobasal media, 1×B27 supplement, 1× Penicillin/streptomycin, 1× L-glutamine).

For live-cell imaging, 35-mm glass-bottom dishes (MatTek) were coated for 1 hour with 0.1 mg/ml poly-D-lysine and then washed 3 times with biology grade water. 2×10^5^ primary cortical cells were plated on the glass coverslip in the center of the dish.

For the 488λ blue LED light exposure of cortical neurons in a time course, 24-well glass bottom black plate (Cellvis, cat#P24-1.5H-N) were used. The plates were pre-coated with 300 μl of 1mg/ml poly-D-lysine for 1 hour at room temperature in the culture hood. Then the plates were washed three times with sterile biology grade water and dry in hood overnight covered in foil. On the day of cell culturing, 2× 10^5^ cortical cells suspended in 200μl plating medium were plated in each well followed by the adding of 1ml feeding medium at 2 hrs after seeding.

For the collection of fresh cell lysate in purpose of biochemistry or Mass spectrometry, 10cm dishes were in use. Briefly, the dishes were coated with poly-D-lysine prior to plating of cortical neurons (4× 10^6^ cells/dish).

For all plate or dish types, the cultures were maintained at 37°C in the incubator with 5% CO_2_ and 95% air. For cell maintenance, ~1/2 volume of feeding media were replaced every 3-4 days until ready to use for experiment on day-11 to day-14.

For cell transduction of lentivirus, at day 2, neurons were transduced with lentivirus vectors at MOI 10 followed by medium change 48 hrs later. And the process were repeated on day5 for a double hit of transduction to increase the transfection efficacy. With this double-hit transduction procedure, the final transfection efficiency will be increased to ~50% while the transfection efficiency with a single time of lentivirus addition was ~30%. For the percentage of neurons response sensitively to light, it will be around 10% to total based on their diverse expression levels.

### Plasmid construction

The mCherry::Cry2 and 4R1N Tau::Cry2 construct were designed by Dr. Brangwynne group. Briefly, sequences for mCherry and Cry2Olig (Addgene) were cloned into the pHR lentiviral backbone to generate the pHR-mCh-Cry2Olig plasmid. DNA fragment encoding human 4R1N full length tau were amplified by PCR using tau cDNA (Addgene). And then the DNA fragment encoding 4R1N tau were inserted into the linearized pHR-mCh-Cry2Olig backbone using In-Fusion Cloning Kit (Takara). The resulting constructs were fully sequenced to confirm the absence of unwanted substitutions.

### Lentivirus production

Lentivirus was prepared as described previously (Araki et al., 2004): HEK293T cells were plated at a concentration of 1×10^6^ cells/well in a 6-well plate. 18 hrs later, cells were transiently co-transfected with PSP (1200 ng), VSV-G (400 ng), and target (400 ng) plasmids using 6 μL FuGene HD (Promega). 72 hrs later, conditioned media was harvested and centrifuged at 1000xg for 5 min to remove dead cells and debris. Supernatant was stored at −80°C until use. For primary neuron transduction, lentivirus was concentrated 10x using lenti-X concentrator (Clontech) with the concentrated pellet being re-suspended in PBS with 25 mM HEPES, pH 7.4.

### Lentivirus transduction

For cell transduction of lentivirus, at day 2, neurons were transduced with lentivirus vectors at MOI 10 followed by medium change 48 hrs later. And the process were repeated on day5 for a double hit of transduction to increase the transfection efficacy. With this double-hit transduction procedure, the final transfection efficiency will be increased to ~60% while the transfection efficiency with a single time of lentivirus addition was ~30%. For the percentage of neurons response sensitively to light, it will be around 10% to total based on their diverse expression levels.

### Knock down gene expression by siRNA

Primary cortical cultures of neurons were transduced with lentivirus coding for Tau:Cry2 (or Cry2Olig) on DIV2 and 4. On DIV6 the cultures were transduced with siRNA HNRNPA2B1 or scrambled control siRNA. On DIV10 the neurons were exposed to 20 or 60 min of 488 λ light, fixed and imaged. The HNRNPA2B1 siRNA (catalog# MBS829521) and control siRNA (catalog# MBS8241404) were purchased from *MyBioSource*. The siRNA oligo duplexes were resuspend and treated in cell culture as directed by the manufacture’s instruction. Briefly, the pre-designed sets of 3 different target-specific siRNA oligo duplexes of mouse HNRNPA2B1 gene were resuspend in each vial with DEPC water. Then the 3 vials of siRNA oligo were combined into one vial. The stock concentration of each oligomer was 100μM. On the day of transfection, 300μl neuronal feeding medium containing 20 pmol of each siRNA oligo was used to completely replace the old medium in each well of the 24-well plate. The olig sequence of each siRNA pool are as follows:

siRNA Negative Control:
UUCUCCGAACGUGUCACGUTT
ACGUGACACGUUCGGAGAATT
siRNA HNRNPA2B1:
HNRNPA2B1 siRNA (Mouse)-A:
CCACCUUAGAGAUUACUUUTT
AAAGUAAUCUCUAAGGUGGTT
HNRNPA2B1 siRNA (Mouse)-B:
CCGAUAGGCAGUCUGGAAATT
UUUCCAGACUGCCUAUCGGTT
HNRNPA2B1 siRNA (Mouse)-C:
CCAGCAGCCUUCUAACUAUTT
AUAGUUAGAAGGCUGCUGGTT

### Live-cell imaging

The live-cell imaging procedure were referenced from precious publications(Shin et al., 2017). Briefly, on day12 to day14 of the primary cortical neurons, prior to imaging, the medium was replaced with imaging medium consisting of 2% FBS in HBSS (Corning cellgro). All live cell imaging was performed using 63× oil immersion objective on a Zeiss LSM880 laser scanning confocal microscope equipped with a temperature stage at 37°C and CO_2_ 5%. For global activation, cells were imaged typically by use of two laser wavelength (488 nm for Cry2Olig activation /560 nm for mCherry imaging). To execute activation protocols with varying activation intervals, the repetitive ON/OFF cycle was applied by varying the length of OFF time (the activation duration, ta, was fixed to 1 s in all measurements). Localized activation experiments were performed using the stimulation setting where the blue laser scans only a designated region of interest.

### Blue light exposure with 488λ LED bulb

Optogenetic gene control utilizes a light-activated protein which transfers light energy from photons to chemical energy, facilitating high-affinity binding for the cryptochrome-2 protein (Cry2Olig). For this research, a Cry2Olig protein which is fused to Tau is engineered to facilitate oligomeric tau formation by inducing strong affinity binding of the Cry2Olig protein. To reach certain threshold energy for Cry2Olig protein, a mounted LED is utilized to reach a light intensity of 2.5nW/mm^2^. Here we utilized the ThorLabs M488L4 LED, which emits 488 nm with a minimum LED power output of 205mW, a forward voltage of 3.8V and a maximum current of 350mA, and the LEDD1B T-Cube LED driver, which regulates at a maximum current of 1.2A.

#### Model and calculation

Two cell culture containers were used for the optogenetic Cry2Olig activation: 100×18mm Petri Dish and 24 well plates by Thermo Fisher. The petri dish has a diameter of 88mm for the inner ring holding the cell and the 24 well plates contains 24 inner rings of 15.6mm. The LED is placed 4 cm above all cell cultures. For 10 cm petri dish, the light activates the whole dish, while for 24-well plate, the light activates half the plate. To activate the Cry2Olig gene, an intensity of 200μW/cm^2^ is required.

The total light power for disks in different diameters is governed by the multiplying two properties (Equation 1):

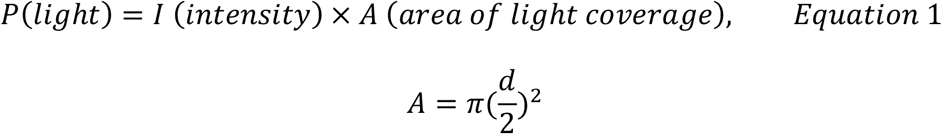

I is the intensity required, 200μW/cm^2^ and d is the diameter of the culture disk and A is the area of the culture disk holding all the cells.

For 10cm petri dish and 24-well plate, the area and light power are shown below (Table 1):

**Table 1.** Light Power Calculation for Culture Disks

|  | 10cm Petri Dish | 24-Well Plate |
| --- | --- | --- |
| Intensity($\mu\text{W}/\text{cm}^2$) | 200 | 200 |
| Height(cm) | 4 | 4 |
| Diameter(mm)/Dimension(mm) | 88 | 85.5x63.9 |
| Area($\text{cm}^2$) | 60.82 | 54.63 |
| Light Power(mW) | 12.2 | 10.9 |

The M488L4 LED, as a photodiode, holds a coefficient for the electron to photon conversion. The maximum output power at a distance of 200mm (20cm) is 240mW. Since the LED is located 4cm above the light cell, by the inverse square law, the intensity increases 25 fold with a maximum irradiance at 4cm of 6W, above our required light power.

Assuming the LED functions linearly and based on the sphere illumines model, the source strength at 20cm is 240mW, governed by the spatial radiation distribution theory with the equation shown below (equation 2). S is the source strength at 4 cm, r is the diameter between the light source and the measured point, and I is the maximum irradiance measured.

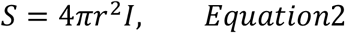

Taken by the experimental data presented in the datasheet. We calculate the coefficient using equation 3 below:

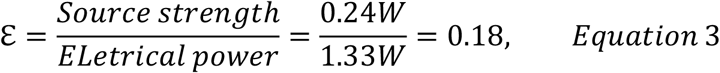

The coefficient for the electron to photon conversion is approximated as 0.945 for ThorLab M488L4LED. Therefore the electrical power is governed by equation 4 below:

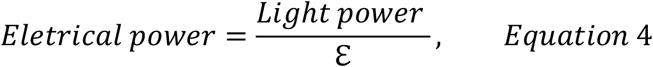

Moreover, the electrical power for a photodiode is governed by the forward voltage and currents shown by equation 5 below:

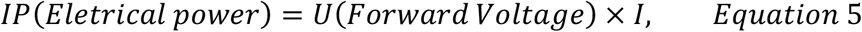

The M470L4 LED has a forward voltage of 3.2 to reach the threshold for proton conversion. Consequently, the power calculations and currents required to activate Cry2Olig protein for different culture disk is shown below in Table 2.

**Table 2.** Electrical Power Calculation for Culture Disks

|  | 10cm Petri Dish | 24-Well Plate |
| --- | --- | --- |
| $\varepsilon$ | 0.18 | 0.18 |
| Light Power(mW) | 12.2 | 10.9 |
| Electrical power (mW) | 67 | 61 |
| Forward Voltage(V) | 3.8 | 3.8 |
| Current(mA) | 17.6 | 16.1 |

The current is controlled by ThorLabs T-Cube LED Driver (LEDD1B) with a maximum current of 1200mA. The designated current is selected by tuning the knob of the system.

### Immuno-fluorescence labeling of fixed primary culture

In the glass bottom 24-well plate with cells, cells were fixed with 0.5 ml 4% PFA/PBS for 15 min followed by being washed 3 times in PBS, 5 min each. In the specific experiments where the m^6^A was to be detected, the cells will be exposed to UV light for 10 minutes before being fixed by PFA. Cells were permeabilized in 0.5ml PBS/0.1% Triton X-100 (PBST) for 15-30 min before being blocked in 0.5ml of 5% BSA - 5% donkey Serum in PBST for 1 hour. And then the cells were incubated in 1° antibody diluted in 5% BSA/PBST overnight at 4°C. Then the cells were washed 3x in PBS-T, 10 min each before being incubated in 2° antibody diluted in 5% BSA/PBST, 2 hrs at RT. All the 2° antibodies were purchased from Thermo Fisher Scientific made in donkey and used for 1:800 dilution in labeling. After 2° antibody, cells were incubated in DAPI diluted 1:10,000 in PBST (5 mg/ml stock solution) for 5 min after first wash. Then cells were washed 2x with PBST, and once of PBS, 10 min each, prior to being covered in Prolong Gold Antifade mounting media with 12mm coverslips. Plates were naturally dried in fume hood (or store at 4°C until ready to dry in fume hood). The primary antibodies used in this study for ICC are as follows: Chicken polyclonal anti-MAP2, 1:250 (Aves Labs, Cat#MAP, RRID: AB_2313549); Goat polyclonal anti-mCherry, 1:300 (MyBioSource, Cat# MBS448057); Rabbit monoclonal anti-Lamin B2, 1:500 (Abcam, Cat# ab151735); Mouse monoclonal anti-Puromycin, clone 12D10, 1: 500 (Millipore Sigma, Cat# MABE343, RRID:AB_2566826); Rabbit monoclonal anti-Cleaved Caspase 3 (Asp175) (5A1E), 1:400 (Cell Signaling Technology, Cat# 9664, RRID:AB_2070042); Mouse monoclonal phospho-Tau (Thr181) antibody AT270, 1:400 (Thermo Fisher Scientific, Cat# MN1050, RRID:AB_223651); Mouse monoclonal phospho-Tau (S262) antibody 12E8, 1:400 (provided by Philip Dolan, Prothena); Rabbit Polyclonal anti-HNRNP A2B1, 1:300 (Thermo Fisher Scientific, Cat# PA534939, RRID:AB_2552288); TIA1 (rabbit, abcam, cat# ab40693, specifically lot# GR3202325-1, 1:400); Monoclonal mouse anti-m^6^A IgG, 1:500 (Synaptic Systems, Cat# 202 111); Mouse monoclonal anti-TOMA2, 1:300 (provided by Rakez Kayed). All the secondary antibodies were purchased from Jackson ImmunoResearch.

### Click-iT™ Plus TUNEL Assay

The Click-iT™ Plus TUNEL Assay kit for in situ apoptosis detection with Alexa Fluor488 dyes was purchased from Invitrogen (Catalog Nos. C10617). The labeling protocol was as instructed by the manufacture’s manual.

### LDH assay

50 μl supernatant were collected as designed time point into a 96-well plate for lactate dehydrogenase (LDH) release assay as manufacture’s protocol (Promega, cat# G1780). Briefly, add 50μl of the CytoTox 96^®^ Reagent to each sample aliquot. The plate was covered with foil to protect it from light, and incubated for 30 minutes at room temperature on shaker. 50μl of Stop Solution was added to each well of the 96-well plate, and the absorbance recorded at 490nm with the plate reader. Each experiment was repeated at least 3 times with triplicate wells each time.

### Surface sensing of translation (SUnSET)

Cells were treated with 10 μg/ml puromycin (Research Products International P33020, resuspended in water) into media for 30mins without or with light for 30mins. Wash twice with ice-cold PBS before lysis. Cell lysate was mechanically lysed in homogenization buffer containing 50 mM Tris (pH 8.0), 274 mM NaCl, 5 mM KCl, with protease inhibitors (Sigma 4693159001), PMSF (1 mM final concentration), and phosphatase inhibitors (Gibco 786-452 and −451). Samples were centrifuged at 4° C at 13,000 g for 15 min, and the supernatant was used for immunoblot. If the visualization of newly synthesized protein by immunolabeling was desired, the cell cultures were fixed by 4% PFA after PBS washing and then the immuno-fluorescence labeling steps were performed as described above. The dilution of puromycin antibody hereby in WB was 1:2000 and in IF labeling 1:1000.

### Immunoprecipitation of mCherry-fusion Proteins

After light exposure, we aspirated the culture medium completely and washed cells with 10ml cold PBS twice, and then the dish was flash-frozen on dry ice. On ice, 800ul homogenization buffer were added in to each 10cm dishes of primary cortical neurons (~4×10^6^ cells) which were transfected with either mCherry::Cry2 or 4R1N Tau::Cry2 lentivirus. The cells were scraped from the bottom of the dish and transfer them into 1.5ml tubes (low binding affinity, cat#z666505). Then the cells were homogenized using cordless motorized tissue grinder with blue plastic pestles, 3 cycles of 30 sec with 10 sec pause. Cell lysate were stored at −80° C for future use.

On the day of mCherry IP, 0.5mM EDTA were added into the cell lysate before it was centrifuged at 20.000x g for 10 min at 4°C. Then the supernatant was transferred to a pre-cooled tube. We prepared the RFP-Trap magnetic agarose beads (Chromotek, Cat# rtma-20) by vortex and pipetted 25 μl bead slurry into 500 μl ice-cold dilution buffer. The beads were magnetically separated until supernatant is clear. We discard the supernatant and repeat wash twice. Then we added the cell lysate to equilibrate RFP-Trap beads following by tumbling end-over end overnight at 4°C. on the second day, we magnetically separated beads until supernatant is clear. After that, we resuspend RFP-Trap beads in 500 μl dilution buffer and repeat wash twice. Then the bound proteins were eluted by adding 50 μl 2×LDS sample buffer followed by boiling at 95 °C for 10 min.

To purify the eluted proteins for the following Nano-LC Mass Spectrometry, we loaded all the samples into a 12% gel and isolated each sample when they went through about 1 cm out of the gel well.

### In-gel digestion and LC-MS analysis

Protein bands were excised and sliced into pieces (2×2mm), then washed with water, 50% Ambic/ACN, and ACN sequentially, and dried. Afterwards, gel pieces were rehydrated with 1ug of trypsin in 100uL of 50mM Ambic with 10% ACN on ice for 30mins, after sufficient Ambic was added to cover the gel piece. Protein in gel were digested at 37 ^o^C overnight followed by the addition of formic acid to 1% in solution. Samples were evaporated to dryness in a vacuum concentrator, and reconstituted in 1% formic acid before LC-MS analysis.

A C18 pre-column (PepMap; 3μm, 100 Å, 75μm × 2cm) hyphenated to a C18 analytical column (RSLC; 2μm, 100 Å, 75μm × 50cm) was used to separate peptides. High performance nanoflow liquid chromatography – Orbitrap tandem mass spectrometry (LC-MS/MS) analyses were completed using an EASY nLC 1200 HPLC system coupled to a Q Exactive HF-X tandem mass spectrometer (Thermo Scientific). Full MS precursor ion profile spectra were collected at a resolution of 60,000 using an automatic gain control of AGC target of 3×10^6^ or a maximum injection time of 50 ms over a scan range of 300 ~1650 m/z. Data-dependent acquisition of high energy collision dissociation (HCD) MS2 fragmentation scans were performed at 15,000 resolution using 27% total normalized collision energy, with dynamic exclusion set to 30 sec.

The acquired spectra was searched by MaxQuant against the UniProt mouse proteome, specifying a fragment ion mass tolerance of 20 ppm, maximum of missed cleavage sites, oxidation as variable modification, and a stringent false discovery rate of 1%. Protein candidates tentatively identified with less than 2 unique peptides were excluded from further analysis.

### Isolation of TBS Soluble (S1p)

The S1p fraction were generated as described previously (Apicco et al., 2018). The cell lysis protocol was done as described in the SUnSET section above. Brain tissue was mechanically lysed in homogenization buffer containing 50 mM Tris (pH 8.0), 274 mM NaCl, 5 mM KCl, with protease inhibitors (Sigma 4693159001), PMSF (1 mM final concentration), and phosphatase inhibitors (Gibco 786-452 and −451). The lysate was centrifuged at 4° C at 13,000 g for 15 min. The resulting supernatant was ultracentrifuged at 28k rpm (29800 x g) in TLA-55 rotor for 20 min at 4 °C. And then we aliquot supernatant to new microtubes as S1 (**TBS-SOLUBLE**). The S1 fraction will be centrifuged @ 60k rpm (150000g) for 60 min at 4 °C (Rotor TLA-55). Then pellet was marked as S1p. S1p and P fraction were then tested by dot blot or immunoblot.

### Immunoprecipitation with PS19 brain tissue

Whole brain cell lysates from 3- and 6-month old PS19 P301S tau transgenic, and WT mice were used for the immunoprecipitation of HNRNPA2B1, HNRNPH, Eif3l and Lamin B2. To prepare the cell lysate, half hemisphere of each brain was lysed in homogenization buffer containing 0.02% NP40. Supernatant from each lysed brain were collected after centrifugation for 15 minutes at 12000g at 4°C and further used for immunoprecipitation.

For immunoprecipitation, 25 ul of completely re-suspended magnetic Dynabeads (Invitrogen, 10003D) were washed 3 times with the homogenization buffer by placing the tubes on the magnet rack to separate the beads from the solution. After washing the beads, 3.5 ug of control rabbit IgG or HNRNPA2/B1 (Invitrogen, PA5-34939), HNRNPH (Bethyl Laboratories, A300-511A-M), Eif3l (Invitrogen, PA531647) and Lamin B2 (Abcam, ab151735) antibodies diluted with total of 200 ul homogenization buffer, were incubated for 1 hour with magnetic Dynabeads. Antibody bound beads were washed 3 times to remove unbound fractions and incubated for 1 hour with 500 ug of respective total brain cell lysates described previously. The brain cell lysates were diluted 4 times with homogenization buffer without NP40, prior to the incubation step. After 1 hour, the beads were washed 2 times with homogenization buffer followed by 4 washes with homogenization buffer without NP40 to avoid the disruption of Tau oligomers. The bound fraction was then eluted by incubating the beads with 20 ul of 200mM Glycine, pH 2.5, for 10 minutes in ice. The elution step was repeated 2 times followed by adding equal volume of 1M Tris pH 10. Eluted fractions from each IP were then loaded in 4-12% SDS PAGE gel with native page running buffer and transferred on nitrocellulose membrane. Each blot was probed with TOC1 antibody to detect oligomeric Tau. The blots were then stripped and reprobed with HNRNPA2B1, HNRNPH, Eif3l and Lamin B2 antibodies respectively (use information shown in the Key Resources Table).

### Immunoblotting

For the detection of high molecular weight 4R1N Tau::Cry2 chimeras, we used a NuPAGE™ 3-8% Tris-Acetate Protein Gels, the samples were prepared homogenization buffer as described above and without detergent. And reducing reagent or boiling steps were also omitted to protect the oligomer structure.

For the detection of TIA1 and LaminB2 in the soluble and insoluble fractions, dot blots were performed using an apparatus obtained from Bio-Rad (Serial number 84BR 30185) when the proteins were transferred to 0.2μm nitrocellulose membranes.

For the puromycin immunoblot to detect the newly synthesized proteins, cell lysate were collected from frozen cultures with RIPA lysis buffer. Reducing and non-reducing protein samples were separated by gel electrophoresis and transferred to 0.2μm nitrocellulose membranes using the Bolt SDS-PAGE system (Life Technologies).

After immunoblot or dot blot, the membranes were blocked in 5% nonfat dry milk (NFDM) in PBS supplemented with 0.025% Tween-20 (PBST) for 1 hour RT, followed by incubation overnight at 4°C in primary antibody diluted in 5% bovine serum albumin/PBST. Primary antibodies used were as follows: Anti-Puromycin Antibody (1:2000, mouse, Millipore, cat# MABE343), Tau13 (1:10,000, Davies Lab, Northwell), TOC1 (1:1000, mouse, Kanaan lab, MSU), Lamin B2 (1:2000, Rabbit, Abcam, cat# ab151735), mCherry antibody (1:500, goat, MyBioSource, cat# MBS448057), Rabbit monoclonal anti-EIF2α (1:500, Cell Signaling Technology, Cat# 5324, RRID:AB_10692650), Rabbit monoclonal phospho-eIF2α (Ser51) (119A11), 1:500 (Cell Signaling Technology, Cat# 3597, RRID: AB_390740); Rabbit Polyclonal anti-eIF3l, 1:500 (Thermo Fisher Scientific, Invitrogen, Cat# PA531647, RRID:AB_2549120), Rabbit Polyclonal anti-HNRNPH, 1:500 (Bethyl Labs, Cat# A300-511A); Rabbit Polyclonal anti-LBR, 1:600 (Proteintech, Cat# 12398-1-AP, RRID:AB_2138334); Rabbit Polyclonal anti-HNRNPA2B1, 1:300 (Thermo Fisher Scientific, Cat# PA534939, RRID:AB_2552288).

### Immuno-fluorescence labeling of fixed brain tissues

For immuno-labeling, selected brain sections of LEnt from bregma −2.8 were washed in PBS for 10 mins and then permeabilized in 0.5ml PBS/0.25% Triton X-100 (PBST). Block tissues in blocking solution supplemented with 5% BSA and 5% normal donkey serum in PBST, 1.5-2 hrs at room temperature (RT). Then sections were incubated in primary antibodies dilute in 5% BSA/PBST and for overnight at 4°C. On the second day, we washed the brain sections 3x in PBST, 15 min each and then incubated the brain sections in 2° antibodies (1:700 for Dylight-/Alexa-conjugated antibodies made in donkey purchased from Thermo Fisher Scientific) in 5% BSA/PBST for 2 hrs at RT. For DAPI nuclei stain, DAPI (1:10,000) was diluted in PBST and incubated with brain sections for 15 min followed by brain section washing 2x with PBST then 1x with PBS, 10 min each. The brain sections were mounted onto microscope glass slides in Prolong gold antifade reagent. Images were captured by Carl Zeiss confocal LSM700. Primary antibodies used for brain tissue labeling were as follows: Mouse monoclonal anti-TOMA2, 1:300 (provided by Rakez Kayed); Rabbit Polyclonal anti-HNRNPA2B1, 1:300 (Thermo Fisher Scientific, Cat# PA534939, RRID:AB_2552288); Rabbit Polyclonal anti-LBR, 1:300 (Proteintech, Cat# 12398-1-AP, RRID:AB_2138334); Rabbit Polyclonal anti-TDP43, 1:300 (Proteintech, Cat# 12892-1-AP, RRID:AB_2200505).

### Proximity Ligation Assay (PLA)

The PLA assay kit were purchased from Millipore Sigma (Cat# DUO92102-1KT) and performed as instructed by the manufacture’s guide and protocol.

### Co-localization analysis by Fiji ImageJ Coloc 2 plugin

The dendritic length measurement of neurons in MAP-2 labeling were quantified using ImageJ plug-ins NeuronJ to trace the MAP2 positive processes(Schmitz et al., 2011). The labeling intensity in immuno-fluorescence were measured by ImageJ. Co-localization of mCherry positive tau oligomers to TIA1 granules in neuronal soma were analyzed with z-stacks images and Pearson coefficient assay by FIJI (ImageJ) coloc2 plug-in. The quantification of cell numbers was done blindly by at least two investigators in the lab.

### Immunoprecipitation of HNRNPA2/B1 from Human AD and Ctrl Brain tissue

Whole brain cell lysates from control and AD human brain tissue were used for the immunoprecipitation of HnRNPA2/B. To prepare the cell lysate, 400 mg of each brain was lysed in lysis buffer containing 50mM Tris, pH 8, 274 mM NaCl, 5mM KCL with 0.02% NP40 supplemented with HALT protease inhibitor and phosphatase inhibitor (ThermoFisher). Supernatant from each lysed brain were collected after centrifugation for 15 minutes at 12000 rcf at 4°C and further used for immunoprecipitation.

For immunoprecipitation, 25 ul of completely re-suspended magnetic Dynabeads (Invitrogen, 10003D) were washed 3 times with the lysis buffer by placing the tubes on the magnet rack to separate the beads from the solution. After washing the beads, 5 μg of control rabbit IgG or HNRNPA2/B1 (Invitrogen, PA5-34939 antibodies diluted with total of 200 μl lysis buffer, were incubated for 1 hour with magnetic Dynabeads. Antibody bound beads were washed 3 times to remove unbound fractions and incubated for 30 min with 5% BSA prepared in lysis buffer to avoid nonspecific binding. The blocking buffer was removed after incubation and the antibody bound beads were incubated for 1 hour with 1 mg of respective total brain cell lysates described previously. The brain cell lysates were diluted 4 times with lysis buffer without NP40, prior to the incubation step. After 1 hour, the beads were washed 2 times with lysis buffer followed by 4 washes with buffer without NP40 to avoid the disruption of Tau oligomers. The bound fraction was then eluted by incubating the beads with 30 μl of 200mM Glycine, pH 2.5, for 10 minutes in ice. The elution step was repeated one more time followed by adding equal volume of 1M Tris pH 10. Eluted fractions from each IP were then loaded in 4-12% SDS PAGE gel with native page running buffer and transferred on nitrocellulose membrane. Each blot was probed with TOC1 antibody to detect oligomeric Tau. The blots were then stripped and reprobed with HNRNPA2/B1 antibody.

### Frozen human tissue samples

| Diagnosis | Age | Sex | Braak Stage | Region |
| --- | --- | --- | --- | --- |
| Control (Assigned ID : 1887) | 60 | M | NA | Frontal cortex |
| Control (Assigned ID : 1886) | 58 | F | NA | Frontal cortex |
| Control (Assigned ID : 1846) | 92 | M | NA | Frontal cortex |
| Control (Assigned ID : 1821) | 92 | M | NA | Frontal cortex |
| Control (Assigned ID : 1703) | 73 | F | NA | Frontal cortex |
| Control (Assigned ID : 1641) | 63 | M | NA | Frontal cortex |
| AD (Assigned ID : 1712) | 87 | F | IV | Frontal cortex |
| AD (Assigned ID : 1620) | 89 | M | IV | Frontal cortex |
| AD (Assigned ID : 1401) | 90 | M | IV | Frontal cortex |
| AD (Assigned ID : 1397) | 88 | M | IV | Frontal cortex |
| AD (Assigned ID : 1345) | 91 | F | IV | Frontal cortex |
| AD (Assigned ID : 1260) | 94 | F | IV | Frontal cortex |

### Bioinformatics analysis of the proteomics data

To adjust for baseline expression, the average value of each protein in each mCherry::Cry2 groups was subtracted from that of corresponding Tau::Cry2 samples at each time point, respectively. Baseline-adjusted protein levels were then normalized to mCherry-Cry2 LFQ protein intensity in each sample. Differential analysis for tau 20min light vs. tau no light, tau 60min light vs. tau 20min light and tau 60min vs. tau no light was performed using the baseline-adjusted, normalized LFQ intensity using linear model with R package limma (Ritchie et al., 2015).

The cutoffs set for volcano plot were fold-change as FC>3 (which is log_2_FC>1.58) and adjust p value p<0.05 for the colored dots in volcano plots, which were labeled as Green FC>3, Red both FC>3 and P<0.05, Blue FC<3 but P<0.05. The cutoff for functional analysis was set as FC>3 and P<0.05. We performed separate GO analyses using the STRING database on proteins which were significantly decreased and those which were significantly increased unique at 20min or 60min of light exposure in comparison to no light group. We also analyzed the Reactome pathways in the functional annotation of STRING database with all the proteins that were significantly changed at 20min or 60min, respectively.

A network analysis was carried out using Cytoscape (V3.7.2). Data from the mCherry IP MS analysis were loaded into Cytoscape and mapped to the STRING protein query database for Mus musculus using the official gene-symbol. The filter for the dataset was established as P<0.05 and (log_2_ FC≥1.58 or log_2_ FC≤-1.58) and a confidence score cut-off was set as 0.8. A protein interaction network based on the STRING interaction score was then formed from the 477 proteins at 20min light group and 372 proteins at 60min group, using the Edge-weighted Spring embedded Layout.

### Statistical analysis

Statistical analyses and figures artwork were performed using GraphPad Prism version 6.00 for Windows with two sided α of 0.05. All group data are expressed as mean ± SEM. Colum means were compared using one-way ANOVA with treatment as the independent variable. And group means were compared using two-way ANOVA with factors on genotype and fractions treatment, respectively. When ANOVA showed a significant difference, pair wise comparisons between group means were examined by Tukey’s, Dunnett or uncorrected Fisher’s LSD multiple comparison test. Significance was defined when *p*< 0.05.

## ACKNOWLEDGEMENTS

We would like to thank Cliff Brangwynne and David Sanders (Princeton) for providing the Cry2Oligo constructs. We thank Peter Davies (Northwell/Hofstra) for provision of the Tau13 antibody, and Philip Dolan (Prothena) for the 12E8 antibody. We would like to acknowledge the Alzheimer Disease Centers of Boston University (NIH grant AG50204517 and P30-AG13846) and Mayo Clinic for donating tissues, as well as the Department of Pathology, Icahn School of Medicine, Mount Sinai Medical Center. We would like to thank the following funding agencies for their support: BW: NIH (AG050471, NS089544, AG056318, AG064932, AG061706), the BrightFocus Foundation and the BU Kilachand Award.

## AUTHOR CONTRIBUTIONS

Conceptualization: B.W. and L.J.; Methodology: L.J., W.L., P.E.A., C.Z., J.K., M.V., S.B., B.F.M., S.L., A.L.C., E.V.V.; Investigation: L.J., W.L., M.V., P.E.A., E.V.V.; Reagents: L.P., R.K., N.K., M.M., J.C., V.E.A. Visualization: L.J., C.Z., H.L., W.L., J.S.; Writing: L.J., B.W.; Editing: B.W., L.J., A.E., W.L., L.P., N.K., P.E.A.; Supervision: B.W., A.E.; Funding Acquisition: B.W, A.E.

